# Amyloidosis and tauopathy progressively disrupt near-critical dynamics in selectively vulnerable circuits

**DOI:** 10.64898/2026.11.11.766020

**Authors:** James N. McGregor, Leandro J. Fosque, Samuel J. Brunwasser, Clayton Farris, Bryan T. Higashikubo, Zachary D. Berriman-Rozen, Halla Elmore, Kylee Kest, Jennifer D. Whitesell, Julie A. Harris, Woodrow L. Shew, David M. Holtzman, Keith B. Hengen

## Abstract

Mechanistically distinct neurodegenerative diseases result in similar cognitive deficits, and pathology is often concentrated in a set of circuits known to be selectively vulnerable. Selective vulnerability is an intrinsic property of a circuit: in APP/PS1 mice, where plaques are global, functional disruptions are confined to circuits within the default mode network. What characterizes the dynamics of vulnerable circuits from those of spared circuits is unknown. We asked whether the distinguishing property is the proximity of population dynamics to criticality, a homeostatic set-point of network activity that determines computational capacity. We recorded single units simultaneously in three selectively vulnerable regions (retrosplenial cortex, anterior cingulate cortex, and CA1 hippocampus) and in primary visual cortex of freely behaving young and old APP/PS1 and wild-type mice. Firing rate and spike-time variability differed by genotype inconsistently across regions and ages. In contrast, three measures from criticality theory each revealed deficits that emerged only with age and only in the vulnerable regions. No measure detected a progressive change in V1. Avalanche scaling is not fixed in neural activity, but varies along a scaling line as the network state changes. As a result, the lines generated by each genotype revealed a constitutive effect of the transgene in young animals, which inverted with age in amyloid-bearing animals, but only in vulnerable circuits. If the destabilization of near-critical dynamics directly reflects a circuit’s functional capacity, the same signature should identify a mechanistically distinct degenerative disease. In a model of tauopathy (P301S/E4), CA1 dynamics were displaced from the control scaling line with the same age-dependent reversal observed in APP/PS1. Our results suggest a network-level principle of neurodegenerative diseases: molecularly unrelated pathologies converge on the loss of near-critical dynamics in selectively vulnerable circuits.

## INTRODUCTION

Brain function arises from the coordinated interactions of hundreds of thousands to billions of neurons. As a result, the most direct account of how mechanisms of degenerative disease translate into behavioral and cognitive symptoms lies at the level of altered neuronal activity. However, identifying the aspects of neuronal activity whose perturbation relates to functional decline is challenging. In some diseases, there are surprisingly few changes in basic features of neuronal activity^1^. In others, there are a multitude of changes^2–4^, even when there is no obvious disruption of cognition and behavior^5,6^. As a thought experiment, consider a circuit whose contribution to cognition is time keeping via a steady oscillation. Neuronal firing rates and the oscillation amplitude could change without computational consequence, whereas any disruption of the oscillation period would impair function. How then can the alterations in network activity that account for disease be identified? Because networks are connected dynamical systems, no single feature of activity can be manipulated experimentally in isolation. The important feature(s) might instead be identified by comparison across time, across circuits, and across diseases. Intuitively, the central feature (analogous to period in the toy example) should depart progressively from normal dynamics, should be evident only in circuits whose function is compromised in disease, and should generalize across mechanistically distinct diseases that produce similar cognitive and behavioral trajectories.

These criteria are not arbitrary; each follows from a property of neurodegenerative disease. First, molecularly unrelated diseases produce syndromes so similar that, even among specialists, a clinical diagnosis of (e.g.) Alzheimer’s disease is frequently overturned at autopsy^7,8^. Nothing about circuit disruption requires such stereotypy. Perturbation of even simple circuits produces diverse, instance-specific failure modes^9,10^. From this perspective, the nearly obligatory progressive cognitive decline in degenerative diseases^11,12^ is remarkable, and instances of disease-driven supranormal function are exceptionally rare^13^. The simplest explanation of this stereotypy is that the same feature of neural dynamics fails across diseases. Second, failure within a brain is anatomically selective. Molecularly unrelated diseases degrade an overlapping set of selectively vulnerable circuits, notably the default mode network (DMN) and hippocampus^14–16^. Third, in human patients, widespread misfolded protein can leave non-DMN circuits functionally intact, at least until late stage disease^17,18^. Disease-relevant dynamics are therefore not simply a byproduct of proximity to pathology; the signature of progressive functional decline is specific to vulnerable circuitry. Together, these properties suggest that a central signature of degenerative disease, should it exist, degrades progressively, is restricted to vulnerable circuitry even when pathology is widespread, and appears across molecular causes.

In Alzheimer’s disease, amyloid plaques accumulate first within the DMN^19,20^ and reach primary sensory cortices only late^21^. Therefore, for most of the disease, pathology and dysfunction occur in the same circuits, and changes in activity driven by amyloid cannot be sorted into those that impair computation and those that do not (period versus amplitude in the toy example). Such sorting requires a brain region that carries amyloid but retains its function alongside vulnerable circuits that are failing. Mouse models of amyloidosis, an analog of early Alzheimer’s disease^22,23^, provide this opportunity. In humans, amyloid is associated with impaired learning^24,25^ and predicts subsequent cognitive decline^26^. APP/PS1 mice similarly develop deficits in learning and memory^27,28^, but do not form neurofibrillary tangles^29^. In both species, the DMN is functionally vulnerable to amyloid and primary sensory cortices are functionally spared^20,30^. However, unlike in humans, amyloid in APP/PS1 mice accumulates throughout the isocortex and hippocampus^31^, rendering vulnerability exclusively functional. Notably, in old APP/PS1 mice, primary visual cortex (V1) can carry a higher plaque density than DMN regions^30^. APP/PS1 mice therefore provide vulnerable and spared circuits that both carry amyloid. Because amyloid impairs cognition in these mice in the absence of tangles, it must drive at least one change in activity that affects computation. A change in V1, where function is spared, is unlikely to bear on functional decline (like amplitude in the toy example). A change that emerges progressively and only in vulnerable circuits would satisfy the first two of our criteria for a dynamical feature central to the impact of disease.

It is possible that a) *all* observed changes in neuronal activity are relevant to cognitive decline, and b) dynamical failures vary by circuit and disease, that is, are not unifiable. Here we test the alternative hypothesis that disease dynamics are unifiable, specifically at the level of a population’s proximity to criticality, a dynamical regime that maximizes information transmission, information storage, dynamic range, and the capacity for adaptive plasticity^32–35^. Like membrane excitability, synaptic strength, and neuronal firing rates^36–40^, proximity to criticality is homeostatically regulated around a set-point^41,42^. This makes proximity to criticality a logical candidate to unify diseases, as it directly constrains computation^34,35^ and neurodegenerative disease can be understood as a progressive challenge to homeostasis that compensation eventually fails to offset^43,44^. Previously, in a mouse model of tauopathy, we found that single-neuron firing was unaffected by disease, whereas proximity to criticality degraded progressively and tracked the extent of tau pathology across animals^1^. However, interpretation of this prior work is limited by its specificity; recordings were restricted to the hippocampus of the P301S/E4 mouse. Further, tau kills neurons^45^, so increased distance from criticality could have been a consequence of degeneration. Studied only in tauopathy, it is thus difficult to parse a dynamical failure of the surviving circuit from the simple loss of cells. In contrast to tau, amyloid causes little neuronal loss^46,47^. If proximity to criticality is the functionally relevant axis onto which both diseases collapse, it should degrade progressively in the vulnerable circuits of APP/PS1 mice, remain intact in V1 despite its higher plaque density, and fail in the same way in tauopathy.

We analyzed chronic recordings of single units made simultaneously in three selectively vulnerable regions, retrosplenial cortex (RSC), anterior cingulate cortex (ACC), and CA1, and in V1 of young and old APP/PS1 and wild-type mice^30^. The transgene is expressed from birth while plaques accumulate with age, so a comparison of young and old animals separates constitutive effects of overexpression from progressive effects of amyloid. Proximity to criticality is not a binary property, and scale invariance can appear in some features of network activity and not others^48–50^. We therefore examined changes in size and duration scaling exponents along a common scaling line^49,51,52^, and two measures of distance to criticality, one computed from avalanches and one from the temporal structure of population activity^41,48^. Single-neuron firing rates and spike-time variability differed by genotype inconsistently across regions and ages. In contrast, in all three analyses, departures from wild-type that emerged with age only appeared in old APP/PS1 mice and only in vulnerable circuits, and V1 showed no progressive change in any of them. In the scaling line analysis, a constitutive effect of the transgene appeared in young animals and inverted with age in vulnerable circuits. McGregor et al.^1^ did not analyze the scaling line, so this signature provided an independent test in tauopathy. In CA1 of P301S/E4 mice, the same age-dependent reversal appeared in the scaling relation. To elucidate the mechanistic changes capable of producing the suite of observed changes, we explored a large space of perturbations in a cortical branching model. Changes in spontaneous activity and in the heterogeneity of input strengths onto neurons were sufficient to reproduce the region-specific changes^53,54^. Together, these results indicate that proximity to criticality meets all three criteria across two molecularly unrelated models of disease, and is a candidate for the feature of neuronal activity whose failure accounts for cognitive decline across circuits and diseases.

## RESULTS

### Simultaneous recording of vulnerable and spared circuits across disease progression in APP/PS1 mice

Homeostatic set-points are defined over hours to days and vary with arousal state^39,42^, so measuring them requires continuous recording across naturally occurring sleep and wake. We therefore reanalyzed chronic recordings of single units in APP/PS1 mice and wild-type (WT) littermates, described previously in the context of interregional communication^30^. Each animal carried four 64-channel tetrode arrays targeting RSC and ACC, two hubs of the DMN, as well as CA1 and V1 (Fig. 1A,B), and lived in an enriched home cage with a littermate while neural activity and video were recorded continuously (Fig. 1C). Every genotype comparison is therefore made between vulnerable and spared circuits recorded simultaneously, in the same animals, across 24 h of free behavior. The transgene is expressed from birth, whereas plaques accumulate with age^46,55^. We recorded young mice (2–4 months; *N* = 4 WT, *N* = 5 APP/PS1), before appreciable plaque deposition, and old mice (12–16 months; *N* = 4 WT, *N* = 4 APP/PS1), during advanced amyloidosis (Fig. 1A). A genotype difference present in young animals is consistent with an effect of the transgene, and one that emerges only in old animals is consistent with progressive amyloid pathology. Single units were classified as regular-spiking (RSU, putative excitatory; *n* = 1765) or fast-spiking (FS, putative inhibitory; *n* = 874) from the bimodal distribution of trough-to-peak durations (Fig. S1A), and arousal state was scored in 4 s epochs (Methods). Because neurons, epochs, and recording blocks are nested within animals, the mouse is the unit of replication (*N* = 17), and every analysis uses a linear mixed-effects model with animal as a random effect and genotype contrasts corrected for multiple comparisons (Methods). All p values reported are false discovery rate corrected.

**Figure 1:**
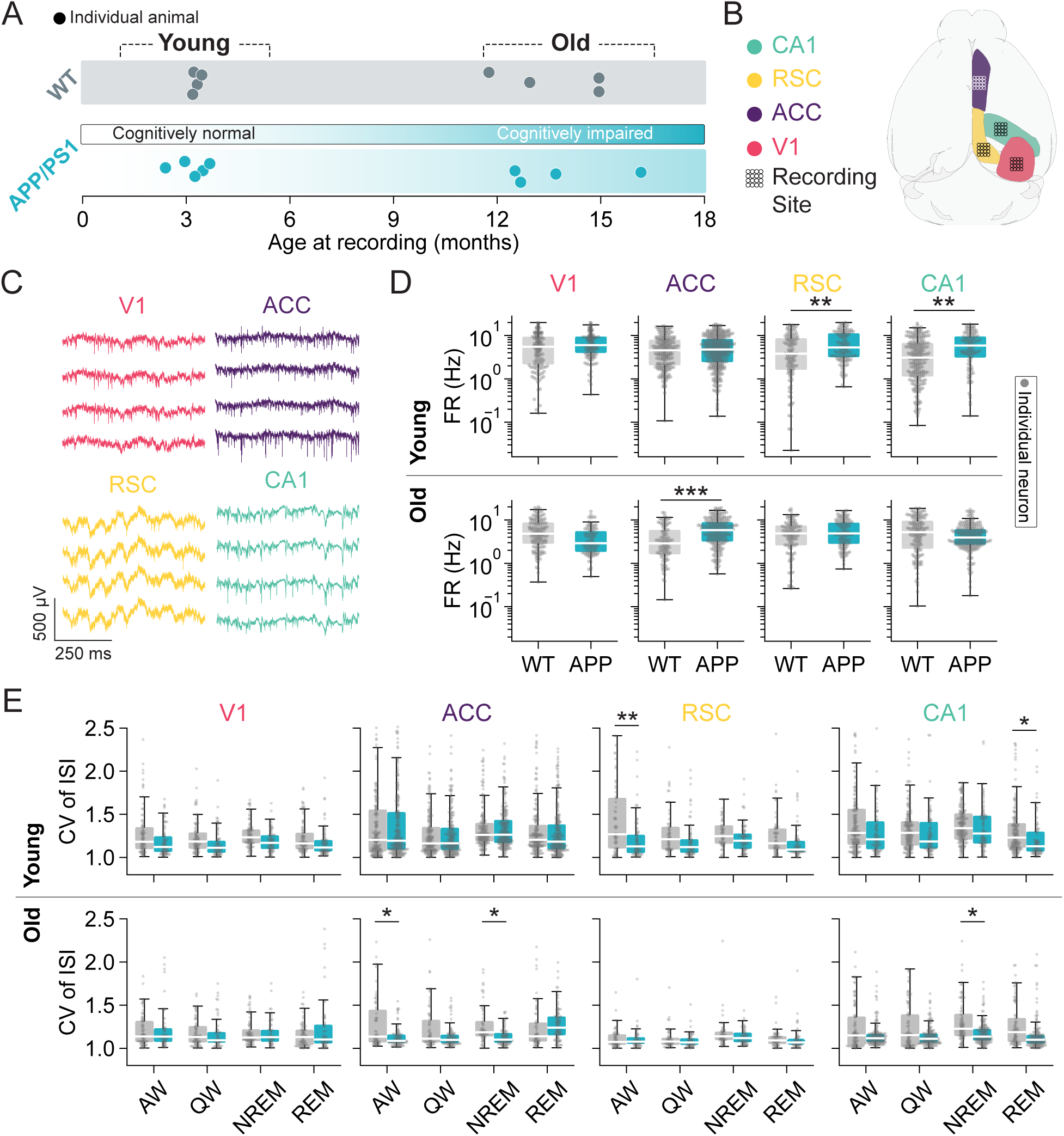
Age-dependent, circuit-specific alterations in single neuron firing rates and spike timing in a mouse model of amyloid accumulation. **A.** Experimental timeline. We recorded neural activity from WT and APP/PS1 mice in two cohorts: young (≤ 4 months; prior to significant plaque accumulation in APP/PS1 mice) and old (≥ 12 months; during late-stage amyloidosis in APP/PS1 mice). The background gradient for APP/PS1 indicates the progressive transition from cognitively normal to cognitively impaired. **B.** 64 ch tetrode bundles were implanted in four brain regions, three associated with higher-order cognition (retrosplenial cortex, RSC; anterior cingulate cortex, ACC; CA1 hippocampus) as well as V1 (primary visual cortex). **C.** Example raw data showing spiking on tetrodes in each of the four recorded regions. **D.** Mean firing rate of regular spiking units (RSUs) as a function of genotype, region, and age group. Firing rate was computed per neuron across the entire recording epoch (spike count divided by time between the neuron’s first and last spike) (* p *<* 0.05, ** p *<* 0.01, *** p *<* 0.001; LMER: log(firing rate) ∼ genotype * age group * region * cell type + sex + prop_NREM_ + prop_REM_ + prop_active wake_ + prop_quiet wake_ + (1 animal), genotype contrasts FDR-corrected). Grey dots indicate individual neuron firing rates. **E.** Coefficient of variation (CV) of the interspike interval (ISI) for RSUs as a function of genotype, region, age group, and brain state (* p *<* 0.05, ** p *<* 0.01; LMER: CV genotype * age group * region * cell type * state + sex + (1 animal/recording epoch/neuron); genotype contrasts FDR-corrected.) Grey dots indicate individual neuron CV values. (AW: active wake; QW: quiet wake; NREM: non-REM sleep; REM: rapid eye movement sleep).

In an independent cohort of 143 APP/PS1 mice, we previously showed that plaque burden in this model is global^30,31^. Plaque density rose by 2.89 log units between 4 and 12–18 months in each recorded region, and in old animals all 43 isocortical regions examined, as well as the hippocampal formation, carried plaques. Among the recorded regions, density was highest in V1 and lowest in CA1 (V1 *>* ACC *>* RSC *>* CA1, all pairwise *p <* 0.0001)^30^. There was no evidence of appreciable neuronal loss. Single-unit yield in old mice did not differ by genotype in any region, and NeuN histology was comparable across genotypes^28,30,56^. As the APP/PS1 mouse is a model of early Alzheimer’s, it is reasonable to expect that altered network dynamics might track the anatomical distribution of plaques, which are expressed globally in this model. Alternatively, if aberrant dynamics are driven by cell death, no changes should occur in APP/PS1. However, if alterations in dynamics reflect the overlap of amyloid and selective vulnerability, they should emerge progressively in RSC, ACC, and CA1 but not V1.

### Amyloid alters single-neuron firing statistics inconsistently across regions and ages

Mean firing rate and the irregularity of spike timing are both under homeostatic control in vivo^38–40^, and tauopathy left both intact in CA1 through end-stage disease^1^, suggesting that these features are not inseparably linked to the degradation of computational capacity in degenerative diseases. However, there is reason to expect a different signature in amyloidosis. Plaques may drive hyperactivity in adjacent neurons^2,57^, and in our prior analysis of these recordings, firing was elevated in RSC and ACC of old APP/PS1 mice^30^. That analysis measured firing within individual arousal-state epochs. Firing-rate homeostasis operates over hours to days^38,39^, and a change in firing within states need not alter the homeostatic set-point. Here we asked whether each neuron’s mean firing rate over 12 h differed by genotype. On the basis of our prior results, we hypothesized that it would be consistently increased in the vulnerable regions of old APP/PS1 animals. If firing rose progressively in every vulnerable region and not in V1, single-neuron activity would meet two of our three criteria in amyloidosis (a progressive departure from normal dynamics and anatomical specificity). In this case, amyloid and tau would disrupt different features of neuronal activity. The null hypothesis is no effect of genotype on firing rate or variability at either age. The remaining possibility is that single-neuron measures change, but inconsistently across regions and ages. Such an outcome would be consistent with effects of the transgene or of amyloid on features of activity that do not account for the loss of function, like amplitude in the toy example.

We modeled log firing rate^58^ with genotype, region, age group, and cell type as interacting fixed effects, sex and the proportion of the interval spent in each arousal state as covariates, and animal ID as a random intercept (Fig. 1D). In old animals, RSU firing rate was elevated in APP/PS1 mice solely in ACC (estimate = 0.634 log units, an approximately 1.9-fold increase, *p* = 3 × 10*^−^*^4^). In young animals, firing rate was elevated in APP/PS1 RSC (estimate = 0.573, *p* = 8 × 10*^−^*^4^) and CA1 (estimate = 0.494, *p* = 0.003). Neither difference was present in old animals. Firing rates in V1 showed no genotype difference at either age. Fast-spiking units showed no genotype difference in any region or age (Fig. S1B). Of the elevations in old animals that we previously reported within arousal-state epochs^30^, the one in ACC persists in the long-term mean and the one in RSC does not. These data show that individual neuron mean firing rates rose progressively in one vulnerable region, but in the other two it was elevated only early in life.

Spike-time irregularity, measured as the coefficient of variation (CV) of the interspike interval within each arousal state, changed in a partly overlapping set of regions (Fig. 1E). In old APP/PS1 mice, RSU CV was reduced in ACC during active wake (estimate = −0.191, *p* = 0.020) and NREM (estimate = −0.174, *p* = 0.036), and in CA1 during NREM (estimate = −0.150, *p* = 0.045). In young APP/PS1 mice, CV was reduced in RSC during active wake (estimate = −0.221, *p* = 0.0067) and in CA1 during REM (estimate = −0.149, *p* = 0.036), and neither difference persisted into old age. Among fast-spiking units, CV was reduced in old APP/PS1 mice in CA1 across all states and in ACC during NREM (Fig. S1C). V1 showed no genotype difference in CV in either cell type, at either age, in any state. Thus, spike timing became more regular in old APP/PS1 mice in ACC and CA1 but not RSC, and RSC and CA1 already differed from wild type in young animals.

Taken together, these data reject both the null hypothesis and the hypothesis of amyloid driving consistent, progressive hyperactivity. Genotype altered single-neuron activity in some of the vulnerable regions but never in V1, despite V1 carrying the highest plaque density of the four recorded regions in old animals. The only progressive increase in mean firing rate was observed in ACC. In RSC and CA1, firing was elevated in young animals and not in old, the inverse of the trajectory expected from plaque-driven hyperactivity and more consistent with a transgene effect. Spike-time irregularity changed in a different subset of regions and ages. Unlike in tauopathy^1^, single-neuron activity does change in amyloidosis, but inconsistently, as predicted for activity features that do not account for the loss of function. We next asked whether population-level features change in a pattern that meets all three criteria, beginning with the avalanche scaling line.

### In old APP/PS1 mice, avalanche scaling departs from wild type in every vulnerable circuit

Activity propagates through neuronal populations in cascades termed neuronal avalanches^49^. An avalanche can be quantified by its size, the number of spikes it contains, and its duration (Fig. 2A). In systems near criticality, the distributions of avalanche size and duration follow power laws, *P* (*S*) ∼ *S^−τS^* and *P* (*D*) ∼ *D^−τD^*, with exponents *τ_S_* and *τ_D_* respectively^41,49^. If mean avalanche size also grows with duration as a power law, ⟨*S*⟩(*T*) ∝ *T^γ^*, the three exponents are not independent and must satisfy the scaling relation *γ* = (*τ_D_*−1)*/*(*τ_S_* −1)^59–61^. Any change in *τ_S_* therefore requires a matching change in *τ_D_*. Such a description of neuronal activity might be expected to be sensitive to pathology; in a grand mal seizure, or any state of synchronous population activity, there is no clear relationship between the number of active neurons and the duration of the event. Empirically, the two exponents vary over time and between recordings, but not independently. Across preparations, species, and states, they fall along a common line in the *τ_S_*–*τ_D_* plane, termed the scaling line^51,52,54^. These exponents can be used to ask three separate questions. First, whether an individual recording satisfies the scaling relation is the basis of the deviation from criticality coefficient (DCC), which was elevated in CA1 of tauopathy mice^1,41^ and to which we return below. Second, where recordings sit along the scaling line has been used to compare states, species, and ages^51,52,54^. Third, the slope of the line describes how much the duration exponent changes for a given change in the size exponent as a circuit moves between states, and could separate conditions such as health and stages of disease.

**Figure 2:**
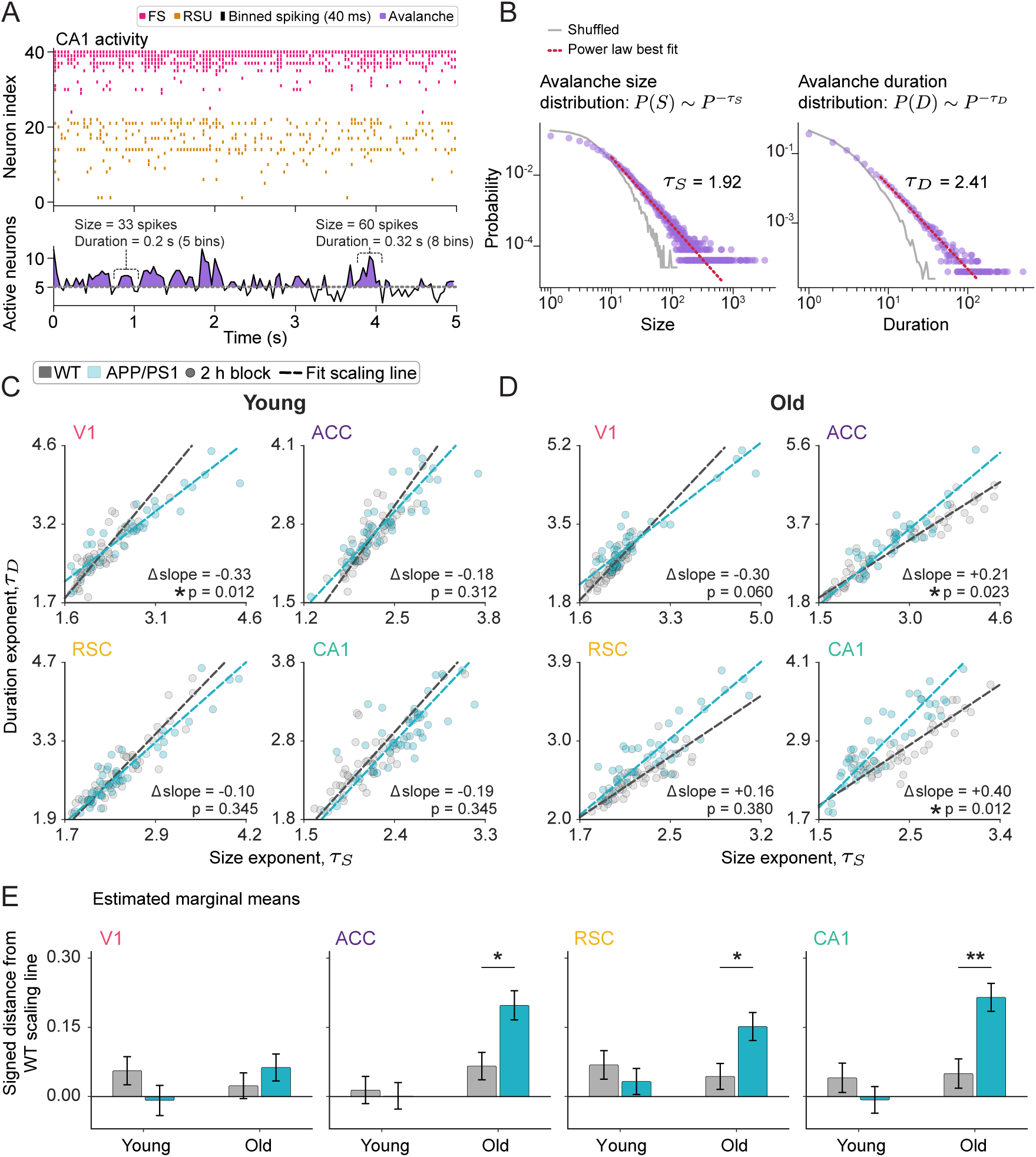
Age-dependent, circuit-specific departure from normal avalanche scaling line in APP/PS1 mice. **A.** Example CA1 population activity. Spiking from RSUs and FSs was binned at 40 ms; avalanches were defined as contiguous periods of population activity above threshold, and each avalanche’s size (total spikes) and duration were quantified. **B.** Example avalanche distributions from a single recording block, fit with power laws to yield the size exponent *τ_S_* and duration exponent *τ_D_* (purple, observed; grey, shuffled; red, power-law fit). **C.** Per-region *τ_S_*–*τ_D_* scatter along the scaling line for young animals. Each point is one 2 h block, colored by genotype (WT, grey; APP/PS1, teal), with a total-least-squares line fit per genotype, region, and age (dashed). Annotations give the difference in scaling line slope between genotypes (Δ slope = APP/PS1 WT) and its significance from the LMER-estimated marginal mean of linear trend interaction, BH-corrected. **D.** Same as **C**, but for old animals. **E.** Signed orthogonal distance of blocks from the WT scaling line. For each region and age, a reference line was fit to WT blocks alone by total least squares, and each block’s signed perpendicular distance from it was computed, so WT centers near zero and positive values indicate displacement above the WT line. Bars, LMER-estimated marginal means SEM (* p *<* 0.05, ** p *<* 0.01, *** p *<* 0.001; LMER: signed distance genotype age group region + splines::ns(N, 3) + sex + (1 animal/recording epoch), Satterthwaite genotype contrasts and Benjamini–Hochberg FDR-corrected).

We reasoned that if avalanche scaling fails selectively in vulnerable regions, then in old APP/PS1 mice the exponents measured in RSC, ACC, and CA1 should depart from wild-type scaling, while those measured in V1 should not. The null hypothesis is that APP/PS1 exponents fall on the wild-type line in every region at every age. A departure could take either of two forms, slope or position. APP/PS1 blocks could occupy a different position along the wild-type line, as different states, species, and ages do^51,52^, or they could fall along a line with a different slope, such that the size and duration exponents covary across states differently in APP/PS1 than WT. Two outcomes would each falsify a specific prediction made in our introduction. First, a departure in V1 as large as in the vulnerable regions would indicate that plaque burden, not circuit identity, determines the effect. Second, a departure present in young animals would indicate an effect of the transgene rather than of progressive pathology.

Spiking from all units within a region, regular-spiking and fast-spiking, was binned at 40 ms and summed. Avalanches were defined as contiguous periods during which population activity exceeded the 35^th^ percentile of its distribution^1,41,42^, and the size (total spikes) and duration of each avalanche were recorded (Fig. 2A). Within each 2 h block, power laws were fit to the size and duration distributions by maximum likelihood, yielding *τ_S_* and *τ_D_*for that block (Fig. 2B). We compared genotypes in three ways: by position along a line fit to both genotypes together, by the slope of a line fit to each genotype separately, and by the signed distance of each block from a line fit to wild-type blocks alone.

Position along a shared line, the axis on which prior work has found conditions to differ^52,54^, did not separate genotypes in any region except RSC in old animals (*p* = 0.048; Fig. S2). In contrast, slopes fit to each genotype separately by total least squares differed between genotypes in CA1 (*p* = 0.012) and ACC (*p* = 0.023) of old animals, and in V1 of young animals (*p* = 0.012). This difference was not significant in old V1 (*p* = 0.060; Fig. 2C,D). These data offer no pattern that completely satisfies our initial predictions of anatomical vulnerability and progression.

We reasoned that slope captures only part of how APP/PS1 blocks could depart from the wild-type scaling line, because blocks could also sit systematically above or below it. To measure this displacement, we computed the signed orthogonal distance of every block from a reference line fit to wild-type data, within each region and age, with the number of recorded units included as a covariate because avalanche exponents depend on population size (Fig. 2E). In young animals, APP/PS1 blocks did not depart from the wild-type line in any region (all *p >* 0.1). In old animals, they departed in ACC (estimate = 0.132, *p* = 0.011), RSC (estimate = 0.108, *p* = 0.033), and CA1 (estimate = 0.165, *p* = 0.003), but not in V1 (estimate = 0.040, *p* = 0.429). Departure from wild-type scaling therefore emerged with age in all three vulnerable regions of APP/PS1 mice and not in V1, despite V1 carrying the highest plaque density, as predicted by selective vulnerability.

Up to this point, we have considered scaling lines as measured in our data. Theory adds an important constraint. A network that receives external input cannot sit exactly at the critical point, and instead operates near it, in a regime termed quasicriticality^53,54^. In a quasicritical network, changes in input move the network toward or away from the critical point and change avalanche exponents, but the exponents continue to approximately satisfy the scaling relation *γ* ≈ (*τ_D_* − 1)*/*(*τ_S_* − 1)^54^. That is to say that input and environment shift a network’s dynamics, but *τ_D_* and *τ_S_* are jointly determined and thus covary along the scaling line. In theory, any line that obeys this relation passes through the point (1, 1)^60^. However, experimentally measured lines rarely do, because thresholding, finite recording time, and error in the fitted exponents shift them up or down. As in prior work^52^, we therefore shifted each genotype’s line vertically so that it passed through (1, 1), leaving its slope unchanged (Fig. 3A,B). Once the wild-type and APP/PS1 lines pass through the same point, they can differ only in slope. Without alignment (Fig. 2E), differences in the vertical offset of the two lines (i.e., their intercept) partly masked differences in slope. After alignment, these differences became apparent. In young animals, APP/PS1 blocks sat below the wild-type line, indicating a shallower slope, in V1 (estimate = −0.416, *p* = 5.8 × 10*^−^*^12^), ACC (estimate = −0.253, *p* = 3.4 × 10*^−^*^7^), and RSC (estimate = −0.168, *p* = 1.8 × 10*^−^*^4^), but not in CA1 (estimate = −0.027, *p* = 0.52; Fig. 3C). This is consistent with a broad (but not universal) transgene effect. This effect persisted in old V1, where APP/PS1 blocks remained below the line (estimate = −0.288, *p* = 6.4×10*^−^*^8^). In stark contrast, all three vulnerable regions reversed direction in old age, such that APP/PS1 blocks sat above the wild-type line. This steeper slope was evident in ACC (estimate = 0.361, *p* = 4.7 × 10*^−^*^10^), RSC (estimate = 0.220, *p* = 6.9 × 10*^−^*^6^), and CA1 (estimate = 0.369, *p* = 4.5 × 10*^−^*^10^). Because both lines are anchored at (1, 1), a given difference in slope produces larger distances in blocks farther from that point. The alignment at (1, 1) therefore magnifies differences between genotypes but does not create them; note that in young CA1, APP/PS1 blocks did not depart from the wild-type line. The free-fit comparison (Fig. 2C,D) tests slope alone, whereas the aligned comparison (Fig. 3C) tests the displacement produced by slope differences, which grows with distance from (1, 1), making the test more sensitive.

**Figure 3:**
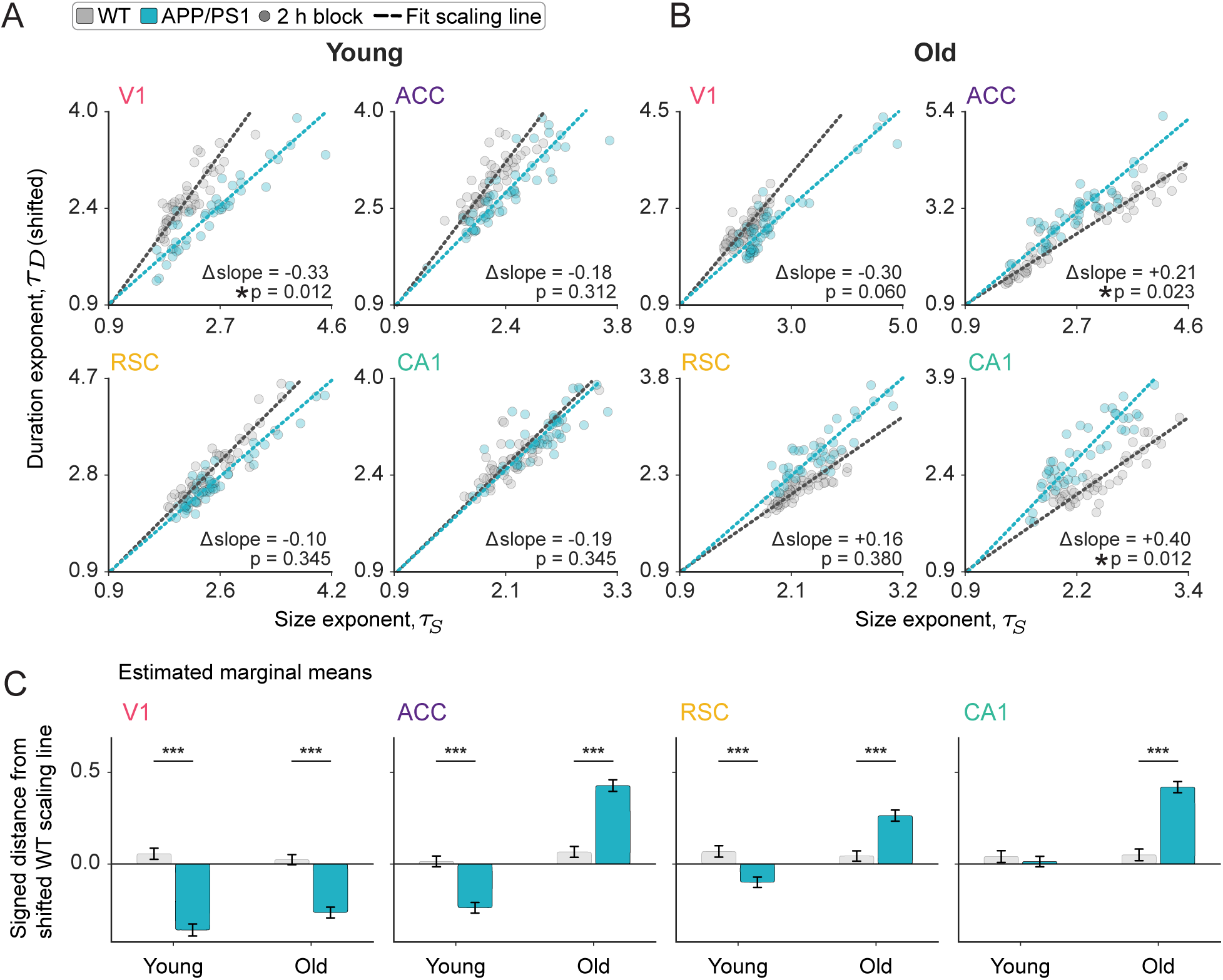
With scaling lines aligned at (1, 1), a transgene effect present in young animals reverses with age in vulnerable circuits. **A.** *τ_S_*–*τ_D_* scatter for young animals in each region. Each point is one 2 h block, colored by genotype (WT, grey; APP/PS1, teal). Each genotype’s duration exponents were shifted so that its scaling line passes through (1, 1), the point through which any line satisfying the scaling relation passes^52,60^. The shift preserves each line’s slope, so slopes and their significance are identical to those in Fig. 2. **B.** Same as **A**, for old animals. **C.** Signed orthogonal distance of each block from the aligned WT line. Because both lines pass through (1, 1), they can differ only in slope, and blocks above the WT line belong to a steeper line. A given difference in slope produces larger distances in blocks farther from (1, 1). Bars, LMER-estimated marginal means SEM; circles, individual-animal medians. Genotype contrasts within each age region combination (* *p <* 0.05, ** *p <* 0.01, *** *p <* 0.001; LMER: signed distance genotype age group region + splines::ns(N, 3) + sex + (1 animal/recording epoch); Satterthwaite degrees of freedom, Benjamini–Hochberg correction).

Plainly, there are two effects of genotype on the scaling line, and they have opposite signs. In young animals, APP/PS1 blocks sat below the wild-type line in every isocortical region, and in V1 this persisted into old age despite extensive amyloid accumulation. This is consistent with constitutive overexpression of the transgene. The second effect appeared only with age and only in vulnerable circuits, where APP/PS1 blocks flipped and sat above the wild-type line, thus reversing the direction of the first effect. This pattern is consistent with an effect of progressive pathology that depends on circuit vulnerability rather than local plaque burden. Surprisingly, neither effect was accompanied by a change in *γ*, which did not differ between genotypes in any region or age (Fig. 4, raw values in Fig. S5). Recall that *γ* describes how avalanches grow in size as their duration increases. There are two plausible explanations of the age, circuit, and genotype-dependent patterns observed here. First, it is possible that the effects are epiphenomenal to this animal model, despite only occurring in selectively vulnerable regions. Alternately, alterations in information propagation captured by changes in the scaling line are an intrinsic feature of cognitive decline. If the latter were true, the same pattern should be observed in the dynamics of a molecularly distinct model of degenerative disease that causes progressive cognitive decline.

**Figure 4:**
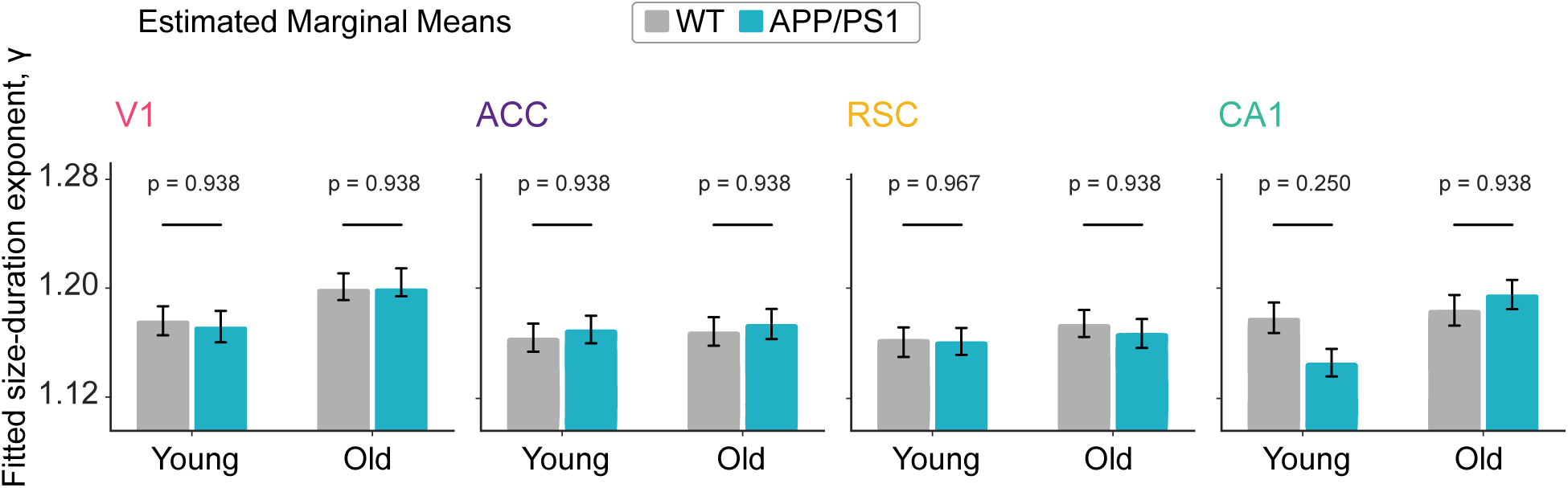
The genotype difference in scaling line slope occurs without a change in the size–duration exponent *γ*. Fitted *γ*, the exponent relating mean avalanche size to duration, *S* (*T*) *T ^γ^*, computed per 2 h recording block and shown as a function of genotype, age group, and brain region. Bars and error bars indicate LMER-estimated marginal means ± SEM. *p*-values above each pair give the genotype contrast within that age region combination. No contrast approached significance (all *p*_FDR_ 0.250). Because the slope differences in Fig. 3 are not accompanied by a shift in *γ*, they reflect a change in how blocks depart from the scaling relation across states rather than a change in the relation itself. (LMER: *γ* genotype age group region + splines::ns(N, 3) + sex + (1 animal/recording epoch), Satterthwaite genotype contrasts and Benjamini–Hochberg FDR-corrected).

We sought to disambiguate these explanations by considering CA1 recordings from our previously published model of tauopathy (P301S/E4)^1^. That study found that proximity to criticality in CA1 degraded as tau pathology progressed, but we did not examine the scaling line, making a direct comparison between the models impossible. To close this gap, we applied the same aligned analysis to young (3–4 months) and old (10–15 months) P301S/E4 mice and littermate controls (23 wild-type and 7 E4 mice; 13 P301S/E4 mice). As in the isocortex of young APP/PS1 mice, the scaling line in young P301S/E4 CA1 was shallower than that of controls (slopes 1.04 vs. 1.24; *p* = 0.018; Fig. 5A). In old animals, it flipped, becoming steeper than controls (slopes 1.56 vs. 1.08; *p <* 0.001; Fig. 5B). P301S/E4 blocks accordingly sat below the control line in young animals (estimate = −0.35, *p* = 4.6 × 10*^−^*^4^) and above it in old animals (estimate = 0.454, *p* = 1.6 × 10*^−^*^4^; Fig. 5C). These significant distances remained in the unshifted scaling line analysis (see Fig. S3). The displacement in old P301S/E4 CA1 was in the same direction as displacement in old APP/PS1 CA1 and of similar magnitude (APP/PS1 estimate = 0.369). Put simply, two molecularly unrelated pathologies steepen the scaling line in CA1 with age, to a similar degree. Unlike young APP/PS1 CA1, young P301S/E4 CA1 already sat below the control line, so in tauopathy CA1 shows the same full reversal seen in APP/PS1 ACC and RSC.

**Figure 5:**
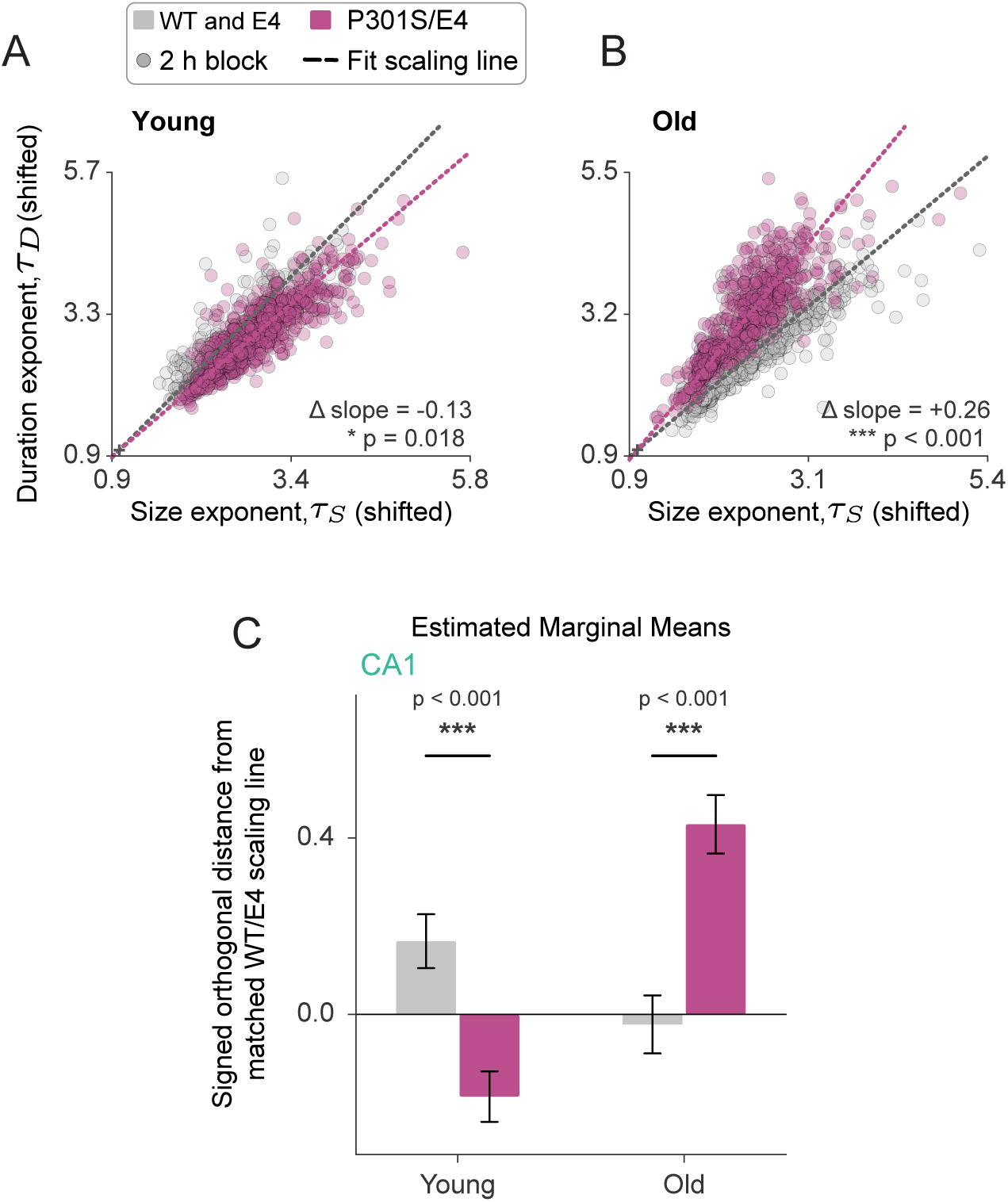
The scaling line departure from wild-type is also present in CA1 in a mouse model of tau-mediated neurodegeneration (P301S/E4). **A.** Per-region *τ_S_*–*τ_D_* scatter on the scaling line from CA1 dynamics for young animals (3-4 mo). Each point is one 2 h block, colored by genotype (control [WT and E4] in grey, and P301S/E4 in magenta). To place both genotypes in a common reference frame, and relate the scaling line slopes to *γ*, each genotype’s duration exponents were shifted so that its scaling line passes through the (1,1)^52^. Total-least-squares lines were then fit per genotype, region, and age (dashed). Annotations give the per-genotype scaling line slopes and the genotype slope difference (same method as in Fig. 2C). **B.** Same as **A**, but for old animals (10-15 mo). **C.** Signed orthogonal distance of CA1 blocks from the control scaling line, measured in this common, aligned frame. Because both genotypes’ lines share (1,1), a block’s perpendicular distance from the control line reflects its departure from the control scaling line independent of where the two lines would otherwise sit. For each age, a reference line was fit to control blocks alone by total least squares, and each block’s signed perpendicular distance from it was computed, so control blocks center near zero and positive values indicate displacement above the control line (same method as Fig. 3).

In disease-vulnerable circuits of old APP/PS1 mice, the scaling line departs from wild-type in the direction opposite to a constitutive effect of the transgene. V1, which is not selectively vulnerable, does not show this age-dependent flip. Tauopathy replicates the age-dependent inversion around the scaling line in CA1. The scaling line describes how avalanche exponents covary across network states, but it does not by itself measure how far a network sits from the critical point. We next asked whether the change in the scaling line corresponds to increased distance from criticality.

### Near-critical dynamics are lost with age in CA1 and RSC of APP/PS1 mice, but not in V1

The deviation from criticality coefficient (DCC) measures how closely the avalanche exponents of a single recording block satisfy the scaling relation^41^. The exponent *γ* can be obtained in two ways, measured directly from how mean avalanche size grows with duration, or predicted from the size and duration exponents as (*τ_D_* −1)*/*(*τ_S_*−1). Near criticality, the two agree giving rise to the relation stated above, *γ* = (*τ_D_*− 1)*/*(*τ_S_*− 1). DCC is the absolute difference between the observed and the predicted (Fig. 6A). Whereas the scaling line compares how exponents covary across blocks, DCC is computed within each block. In our prior work, DCC rose in CA1 as tauopathy progressed^1^. If the change in the scaling line in APP/PS1 mice reflects increased distance from criticality, DCC should be elevated in vulnerable circuits of old APP/PS1 mice, but neither in V1 nor in young animals. The null hypothesis is that genotype has no effect on DCC in any region at either age.

**Figure 6:**
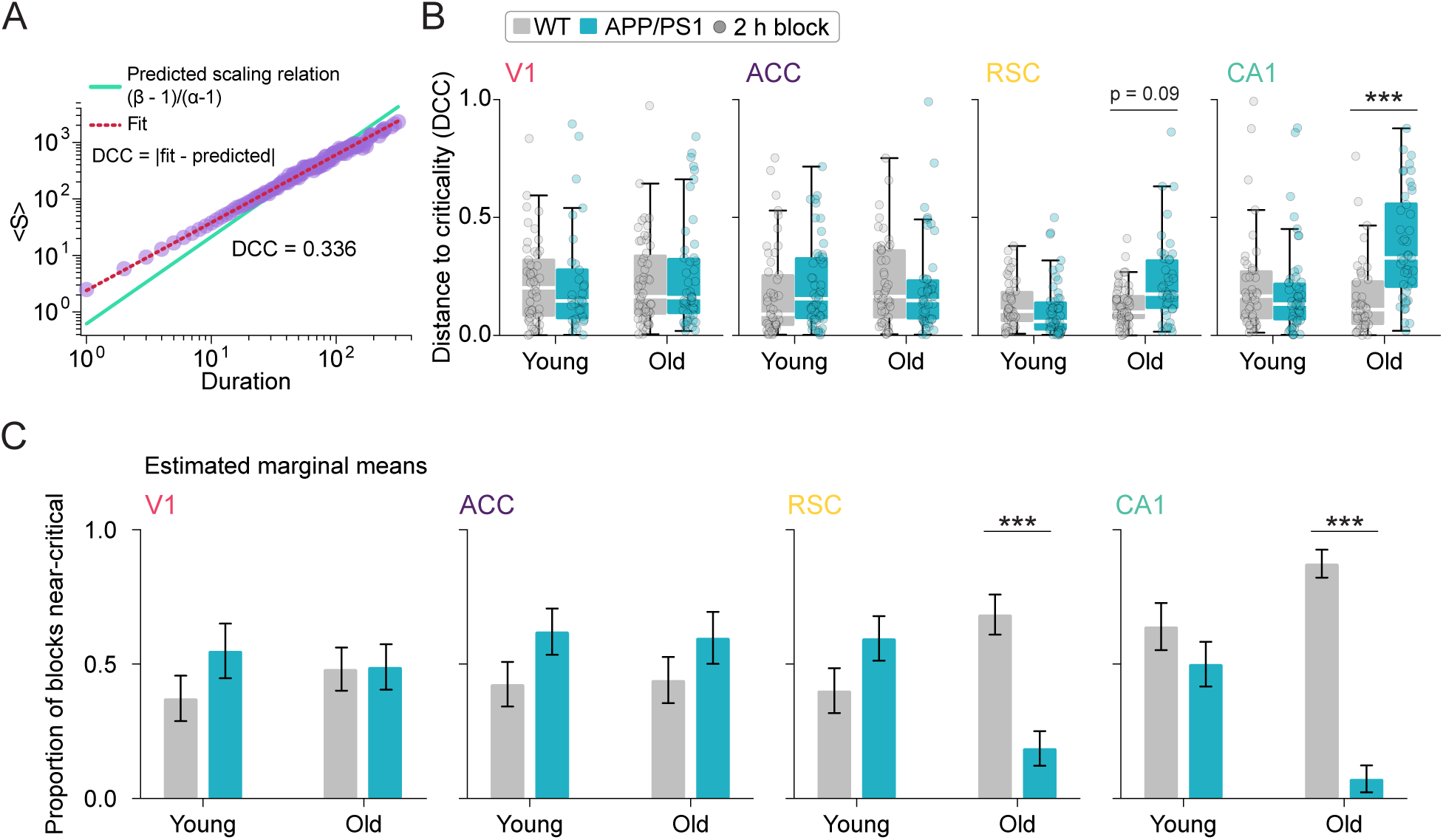
Age-dependent, circuit-specific disruption of near-critical neural dynamics in APP/PS1 mice. **A.** Schematic of the deviation from criticality coefficient (DCC). From the avalanche size and duration exponents (Fig. 3), the scaling relation predicted by criticality, (*τ_D_* 1)*/*(*τ_S_* 1) (cyan), was compared with the empirically fitted relationship between avalanche size and duration (red). DCC was defined as the absolute difference between the predicted and fitted exponents, with values closer to zero indicating dynamics closer to criticality. **B.** DCC across 2 h recording blocks as a function of genotype, age group, and brain region. The shaded yellow region denotes DCC 0.3 (near-critical). Each dot represents an individual 2 h block; boxplots show the distribution across blocks. (* p *<* 0.05, ** p *<* 0.01, *** p *<* 0.001; LMER: DCC genotype age group region + splines::ns(N, 3) + sex + (1 animal/recording epoch); genotype contrasts Benjamini–Hochberg FDR-corrected.) **C.** Proportion of recording blocks classified as near-critical (DCC 0.3) as a function of genotype, age group, and brain region. Circles indicate the observed proportion of near-critical blocks for individual animals; bars and error bars indicate binomial GLMM-estimated proportions SEM. (* p *<* 0.05, ** p *<* 0.01, *** p *<* 0.001; binomial GLMM: near-critical genotype age group region + splines::ns(N, 3) + sex + (1 animal:recording epoch); genotype contrasts Benjamini–Hochberg FDR-corrected.)

We modeled DCC with the same mixed-effects structure used for the scaling line (Fig. 6B). In young animals, DCC did not differ between genotypes in any region (all *p >* 0.2). In old animals, DCC was elevated in APP/PS1 CA1 (estimate = 0.200, *p* = 1.7 × 10*^−^*^4^), with a smaller increase in RSC that did not reach significance (estimate = 0.096, *p* = 0.091). DCC in old animals did not differ in V1 (estimate = 0.058, *p* = 0.30) or in ACC (estimate = −0.013, *p* = 0.86), although the ACC scaling line departed from that of wild-type mice in the same animals. This suggests that DCC detected a progressive departure from criticality in CA1, but not the change in ACC revealed by the scaling line.

DCC does not take into account whether avalanche distributions are well described by power laws. We therefore also classified each block as near-critical if its size and duration distributions both passed power-law tests and DCC was at most 0.3^1^, and modeled each block’s near-critical status (yes or no) with a binomial generalized linear mixed model of the same structure, which estimates the proportion of near-critical blocks in each genotype, age, and region (Fig. 6C). In old APP/PS1 mice, near-critical blocks were less prevalent in both RSC and CA1. In RSC, the estimated proportion of near-critical blocks fell from 0.64 in wild-type mice to 0.18 in APP/PS1 mice (odds ratio = 8.3, *p* = 3.3 × 10*^−^*^4^), and in CA1 it fell from 0.82 to 0.06 (odds ratio = 70.0, *p* = 8.6 × 10*^−^*^6^). Near-critical prevalence did not differ by genotype in V1 or ACC at either age (all *p >* 0.2), or in any region of young animals (all *p >* 0.1). Thus, most blocks in old wild-type RSC and CA1 were near-critical, and most blocks in old APP/PS1 RSC and CA1 were not.

Our avalanche analyses pooled regular-spiking and fast-spiking units, whereas prior studies of criticality in cortex used regular-spiking units alone^41,42^. Restricted to regular-spiking units, DCC remained elevated in old APP/PS1 CA1 (estimate = 0.182, *p* = 9.3×10*^−^*^4^), and near-critical blocks remained less prevalent (odds ratio = 7.0, *p* = 0.014; Fig. S6). In RSC, neither measure differed between genotypes when restricted to regular-spiking units (DCC, estimate = −0.011, *p* = 0.90; near-critical prevalence, *p* = 0.47). The loss of near-critical dynamics in CA1 is therefore present in putative excitatory neurons alone, whereas in RSC it appears to depend on the inclusion of fast-spiking units.

Taken together, DCC and the prevalence of near-critical blocks suggest that the dynamics of old APP/PS1 CA1 and RSC are farther from criticality than those of wild-type mice. This did not hold in V1 or in young animals. These are the same measures that detected the impact of tauopathy in CA1^1^. Notably, neither measure of distance from criticality detected a change in ACC, despite the fact that the scaling line departed from wild-type mice and mean firing rate rose.

Sleep disruption accompanies preclinical Alzheimer’s disease^62^, and in mice sleep loss increases amyloid and tau^63,64^, consistent with a bidirectional relationship in which disrupted sleep promotes pathology^65,66^. Sleep also restores cortical networks toward criticality after waking activity moves them away^42^, and in tauopathy the loss of criticality was most pronounced in wake^1^. Because each 2 h block mixes arousal states, a deficit confined to one state could be diluted in measures computed over whole blocks. Moreover, criticality is not only a property of avalanches. It can also be measured from the temporal structure of population activity, and scale invariance can appear in one form and not the other^48^. We therefore asked whether distance to criticality measured from temporal structure, within single sleep and wake bouts, would reveal deficits, in ACC or elsewhere, that avalanche statistics miss.

### During wake, temporal criticality is disrupted in ACC and RSC of old APP/PS1 mice

A recently developed measure of the distance to temporal criticality (*d*_2_) is computed from the temporal structure of summed population activity^35,48^. Within each 3-minute window, population activity binned at 15 ms was fit with an autoregressive model, which predicts each time bin from the bins that preceded it. Near criticality, fluctuations in activity persist, the autoregressive coefficients sum to nearly one, and *d*_2_ approaches zero. Away from criticality, the influence of past activity decays more quickly, and *d*_2_ increases toward its value for white noise (Fig. 7A,B; Methods). Because a 3-minute window can be assigned to a single arousal state, *d*_2_ can be compared between genotypes separately in wake and in sleep. We modeled window-level *d*_2_ with the same mixed-effects structure used above, with arousal state as an additional interacting factor (Fig. 7C). If the vulnerable circuits of old APP/PS1 mice operate farther from criticality, *d*_2_ should be elevated there, in wake, in sleep, or in both, and not in V1 or in young animals.

**Figure 7:**
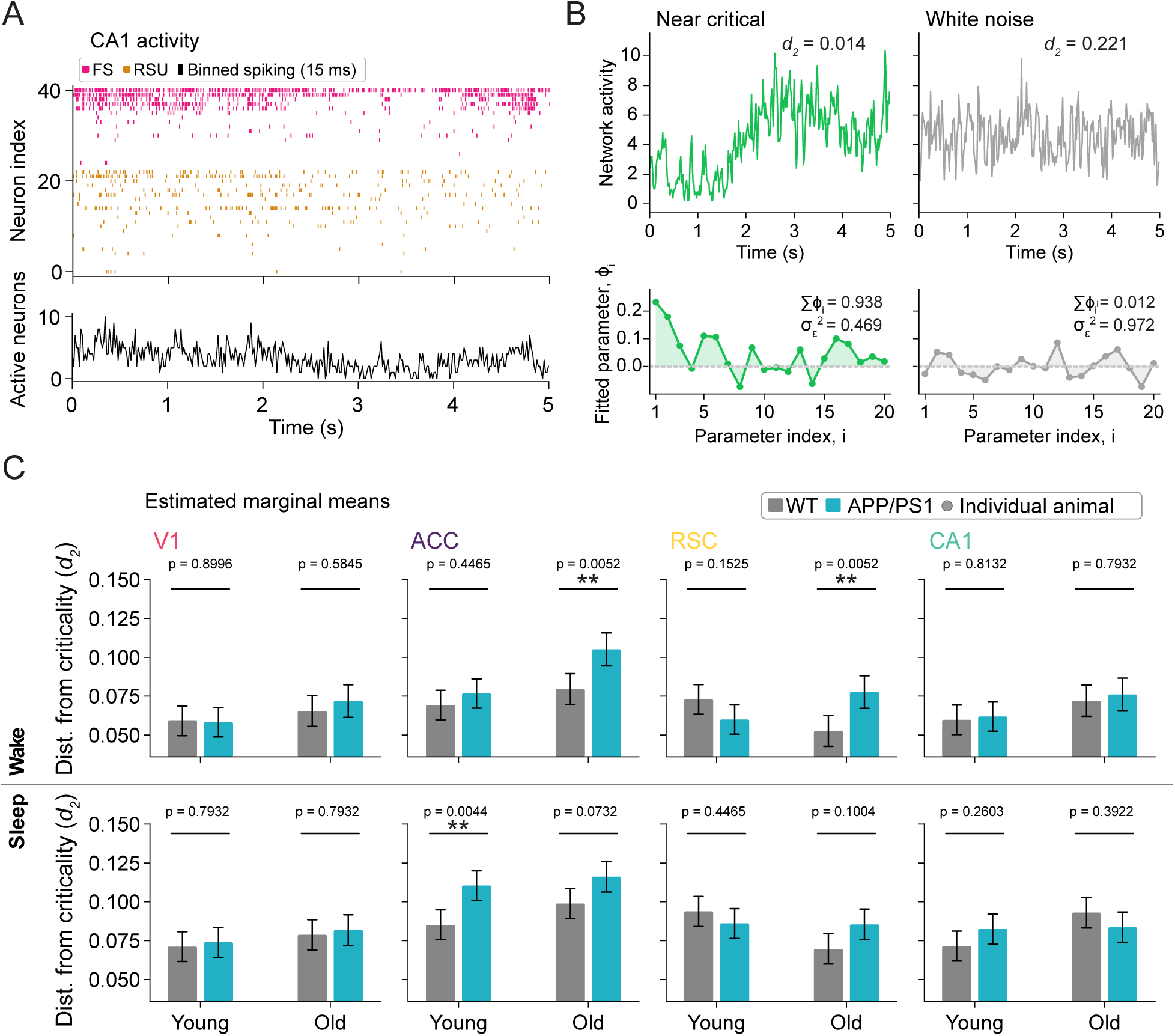
A distinct, state-resolved measure of temporal scale-invariance reveals age-, circuit-, and state-dependent disruptions of criticality in APP/PS1 mice. **A.** Example CA1 population activity. Spiking from regular-spiking units (RSUs) and fast-spiking units (FSs) was binned at 15 ms resolution and summed across the population to yield a network activity time series (bottom). **B.** Schematic of the distance-to-criticality metric (*d*_2_). An autoregressive (AR) model of order *p* = 20 was fit to the network activity time series, predicting each time bin as a weighted sum of the preceding *p* bins with weights *φ_i_* estimated by the Yule–Walker method; 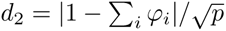 quantifies the distance from criticality. Left: a near-critical 5 s window, with diverse, self-sustaining fluctuations (top); the fitted AR weights sum to nearly one (Σ*_i_ φ_i_* = 0.938), so *d*_2_ approaches zero (bottom). Right: a white-noise window, in which the past of 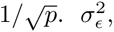 residual noise variance. Values closer to zero indicate dynamics closer to the critical point. **C.** LMER-estimated *d*_2_ as a function of genotype, age group, and brain region, shown separately for wake (AW + QW; top) and sleep (NREM + REM; bottom) states. Circles indicate individual-animal median *d*_2_; bars and error bars indicate LMER-estimated marginal means ± SEM. (* p *<* 0.05, ** p *<* 0.01, *** p *<* 0.001; LMER: *d*_2_∼ genotype ∗ age group ∗ region ∗ state + splines::ns(N, 3) + sex + (1— animal/recording epoch); genotype contrasts Benjamini–Hochberg FDR-corrected across 16 state × age × region comparisons).

In young animals, *d*_2_ did not differ between genotypes in any region or state, with one exception: it was elevated in the ACC of young APP/PS1 mice during sleep (estimate = 0.025, *p* = 0.0044), an effect absent in old animals (estimate = 0.017, *p* = 0.073). In old animals, *d*_2_ during wake was elevated in APP/PS1 ACC (estimate = 0.026, *p* = 0.0052) and RSC (estimate = 0.025, *p* = 0.0052). *d*_2_ did not differ between genotypes in V1 or CA1 in either state (all *p >* 0.05). This measure of temporal criticality detected the deficit in ACC that avalanche statistics missed, but only during wake, and did not detect the deficit in CA1.

Avalanche statistics can also be resolved by state, at coarser resolution, by separating 2 h blocks into those spent mostly asleep (sleep-dense, *>* 50% of the block) and mostly awake (wake-dense)^1^ (Fig. S7). In CA1, DCC was elevated in old APP/PS1 mice in both sleep-dense (estimate = 0.160, *p* = 0.006) and wake-dense blocks (estimate = 0.222, *p* = 0.046), and near-critical blocks were less prevalent in both (sleep-dense, odds ratio = 71.4, *p* = 0.002; wake-dense, odds ratio = 169.7, *p* = 0.023). In RSC, near-critical blocks were less prevalent only in sleep-dense blocks (odds ratio = 9.4, *p* = 0.003; DCC, estimate = 0.098, *p* = 0.107), and neither measure differed between genotypes in wake-dense blocks (DCC, *p* = 0.91; near-critical prevalence, *p* = 0.38). Neither measure differed in V1 or ACC in either state (all *p >* 0.19). The three vulnerable circuits therefore departed from criticality in different states and in different forms of criticality. CA1 lost avalanche criticality in both sleep and wake, with no change in temporal criticality. RSC lost avalanche criticality during sleep, consistent with impaired sleep-dependent restoration of criticality^42^, and temporal criticality during wake. ACC lost temporal criticality during wake alone. Put simply, every vulnerable circuit of old APP/PS1 mice departed from criticality by at least one measure, and V1 did not depart by any.

### In a branching network, concentrating each node’s input on fewer partners reproduces the steeper scaling lines of vulnerable circuits

In the vulnerable circuits of old APP/PS1 mice, the slope of the scaling line changed while *γ* did not. Under the scaling relation, the slope of a line through (1, 1) equals *γ*^60^. As a result, two genotypes with the same *γ* but different slopes must differ in how closely their exponents satisfy the scaling relation. Despite this departure, APP/PS1 blocks continued to fall along a line. Quasicriticality^54^ explains the scaling line as the path of a network whose exponents approximately satisfy the scaling relation as external input moves it toward or away from the critical point^54^. Current work in this theory, however, have not shown if the *τ_S_* and *τ_D_* exponents would remain coupled along a line once the relation is violated. The mechanistic change in network organization capable of producing a new line at constant *γ* is not known. We therefore asked which, if any, changes to a cortical branching model network are sufficient to reproduce the observed, region-specific changes in activity and scaling line slope.

We simulated the cortical branching model, a probabilistic network that can be tuned toward and away from criticality and has been used to explain quasicritical dynamics along the scaling line^53,54^. Each of 128 nodes receives input from 10 others, and the probability that an input activates a node is set by three parameters (Methods). *κ* sets the overall strength of interactions, and, in the absence of external input, the proximity to criticality. *B* sets how unevenly that strength is distributed across a node’s inputs. At *B* = 0 all inputs are equally effective, and as *B* increases, activation depends increasingly on the few strongest inputs. *p_s_* sets the rate of spontaneous activation. For all the simulations, we fixed the number of incoming neighbors to *k_in_* = 10, and refractory period to *τ_r_* = 1. Our main goal was to identify the parameter sets under which the scaling line changed slope relative to a reference line representing wild-type mice, as it did in APP/PS1 mice. For the reference model, which generated the reference scaling line, we chose a slightly subcritical network to allow for exploration without extending to supercritical regimes (*κ* = 1.0062, *B* = 1.5, *p_s_* = 3.16 × 10*^−^*^4^). Then, we searched a grid of values of *κ*, *B*, and *p_s_* (Methods). To match the simulations to the data, we selected the parameter combination for each region whose differences from the reference matched the sign, and then approximately the magnitude, of four differences between old APP/PS1 and wild-type mice: mean population activity, its variance, its coefficient of variation, and the slope of the scaling line (Fig. 8B,C).

**Figure 8:**
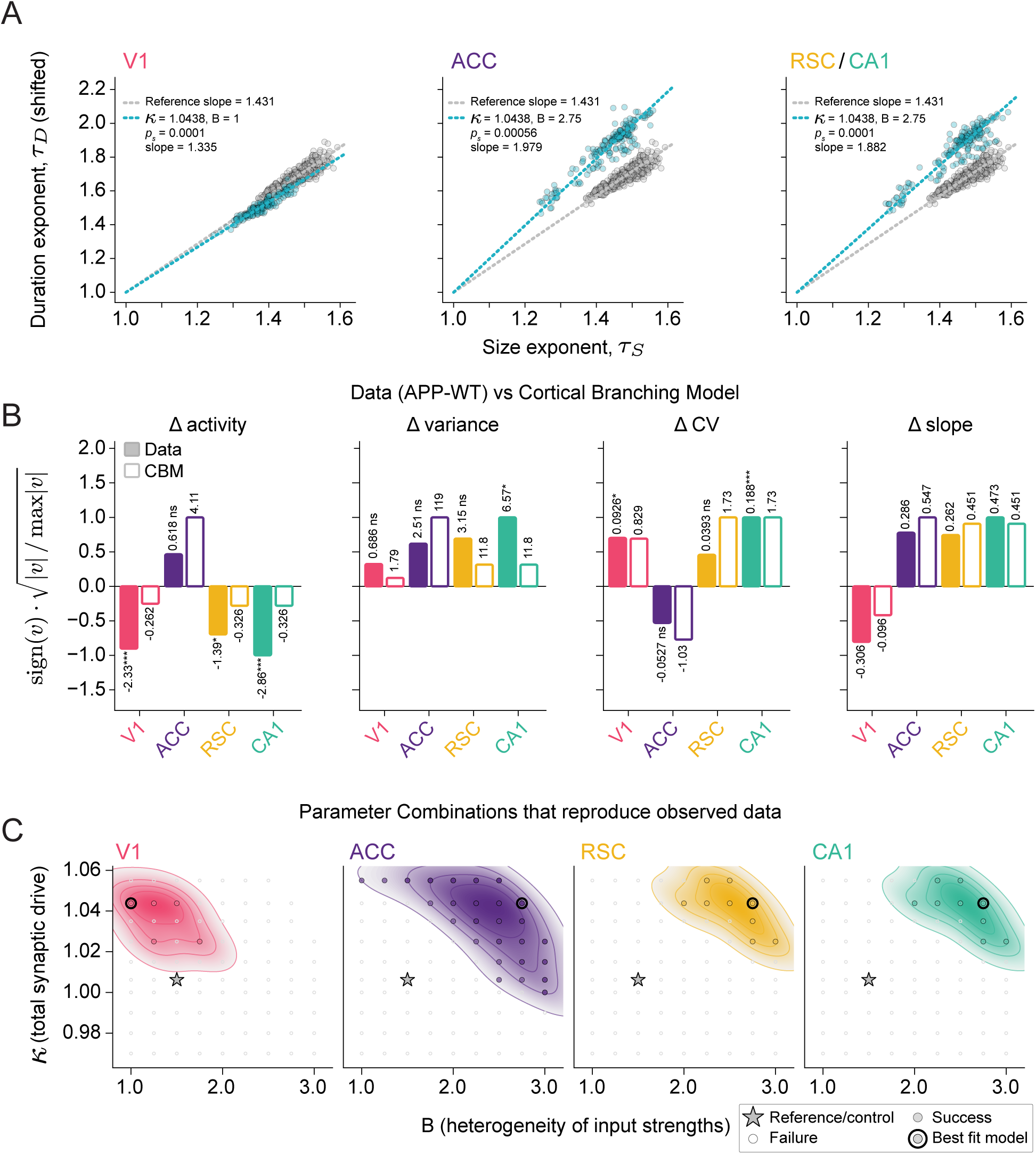
In a cortical branching model, concentrating each node’s input on fewer partners reproduces the steeper scaling lines of vulnerable circuits. **A.** Scaling lines from simulations of the cortical branching model, shifted along the duration-exponent axis to pass through (1, 1) as in Fig. 3. Grey, reference model representing WT (*κ* = 1.0062, *B* = 1.5, *p_s_* = 3.16 10*^−^*^4^); teal, best-fitting model for each region, with parameters and slope annotated. All region models settled at the same *κ* (1.0438). RSC and CA1 matched in the sign of all four measures in **B** and converged on one solution. **B.** The four measures used for the parameter search: the mean, variance, and coefficient of variation of population activity, and the slope of the scaling line. Δ is old APP/PS1 − WT<u>in the data</u> (filled) and region model − reference in the model (open). Bar heights are rescaled as sign(*v*) 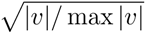 to lie within [−1, 1]; raw values are printed above each bar. Significance on the data bars is from LMER genotype contrasts for each measure (ns, not significant; * *p <* 0.05, ** *p <* 0.01, *** *p <* 0.001). **C.** Parameter combinations searched. Filled circles, combinations matching the sign of all four measures for at least one value of *p_s_*; open circles, combinations that did not; contours, density of matching combinations. Star, reference model; bold circle, best-fitting model in **A**. Sweep ranges are given in Methods.

In the model, as in the data, networks remained on a scaling line while violating the scaling relation (Fig. 8A). All four region models settled at the same value of *κ* (1.0438), higher than that of the reference network, although *κ* was free to vary across the search (Methods). The models for the vulnerable circuits differed from the model for V1 in how input strength was distributed. In the models for ACC and for RSC and CA1, *B* rose from 1.5 to 2.75, so that each node was activated mainly by a few of its inputs, and the scaling line steepened (slopes of 1.979 and 1.882, respectively). In the V1 model, *B* fell to 1, and the scaling line became shallower than that of the reference (slope of 1.335), as in old APP/PS1 V1. Spontaneous activation distinguished ACC from the other vulnerable circuits. The ACC model required more spontaneous activation than the reference (*p_s_* = 5.6 × 10*^−^*^4^), whereas the models for V1 and for RSC and CA1 required less (*p_s_* = 1 × 10*^−^*^4^), consistent with ACC being the only region in which mean firing rate rose in old APP/PS1 mice. In summary, the model reproduces the steeper scaling lines of vulnerable circuits by concentrating each node’s input on fewer partners, without any difference from V1 in proximity to its wild-type reference.

## DISCUSSION

Here we asked whether the circuits that lose function in neurodegenerative disease are unified by a common deficit in neuronal activity. Should such a universal feature exist, we reasoned that it must depart progressively from normal dynamics, should do so only in vulnerable circuits, and should do so across mechanistically distinct diseases. To address this question over time (progression) and anatomy (resilient vs vulnerable) we examined multi-region recordings of single unit ensembles in a mouse model of amyloidosis (APP/PS1^30^). To address the question of mechanistic generalizability, we also analyzed data from CA1 recordings in a mouse model of tauopathy (P301S/E4)^1^. In comparison to WT littermates in each disease cohort, we observed likely transgene effects and sporadic alterations in basic features of neuronal activity, such as firing rate. Consistent with recent hypotheses that degenerative diseases should be defined by the failure of key homeostatic set points in neuronal activity^44^, the only feature that changed progressively with age but only in vulnerable regions and in both disease models was proximity to criticality, a dynamical regime that emerges from tuning to a boundary in parameter space (such as the temperature separating liquid and gaseous phases of matter^34^). General computational capacity is maximized at criticality^32–34,49,67,68^. Thus, its disruption in all combinations of disease-relevant age, anatomy, and mechanism is consistent with the stereotypical decline in functional capacity that defines symptomatic progression across diseases. Intriguingly, our analyses suggest that clear identification of disease-related circuit dynamics, particularly as they pertain to proximity to criticality, may benefit from the application of complementary quantitative tools.

Because causal perturbation of neuronal activity, via, e.g., optogenetic control^69^, invariably alters many aspects of the brain’s dynamics^70^, it would be challenging to test the possibility of a common computational impact of disease via an intervention-based approach. Here we present a theory-driven, comparative, analysis-based approach to the same problem. We identified candidate features by comparison across ages, circuits, and diseases. Crucial to our ability to parse potentially relevant from orthogonal changes (i.e., unrelated to function) is a set of conditions in which pathology and functional decline separate. In Alzheimer’s disease, amyloid accumulates first in the DMN and reaches primary sensory cortices late^19,21^, so for most of the disease timecourse, the circuits that carry amyloid and the circuits that are failing are the same. This does not allow for the separate observation of functionally-compromised circuits and intact circuits that carry molecular pathology. Without this separability, there is no capacity to identify the basis of period and amplitude (in our toy example). Fortunately, in APP/PS1 mice, amyloid is global, and V1 carries amyloid while retaining its function. Note that although retinal changes have been reported in APPswe/PS1dE9 mice^71,72^, pattern visual evoked potentials recorded from awake, aged animals are normal^73^, and baseline visual cortical responses and functional organization are preserved^74^, indicating that visual input reaches and drives V1 largely normally in this line. In contrast, cognitive function and behavior that arise from selectively vulnerable regions are disrupted^75–77^. As a result, comparison of dynamics across regions in the APP/PS1 mouse allows for the rejection of any features that appear in regions that are functionally intact but bearing amyloid (V1, in this case). By definition, those changes are not sufficient to account for functional decline.

Similarly, comparing young and old animals separates constitutive effects of the transgene from progressive effects of pathology. Meeting the three criteria makes proximity to criticality a candidate for the feature whose failure accounts for cognitive decline. Despite its alignment with prior studies of function, our data do not establish causality. Establishing causation would require restoring proximity to criticality in vulnerable circuits and asking whether learning and memory improve as a result.

Proximity to criticality maximizes information transmission, information storage, dynamic range, and the capacity for adaptive plasticity^32–35^, and it is homeostatically regulated around a set-point^41,42^. In old APP/PS1 mice, every vulnerable circuit departed from criticality by at least one measure. In contrast, V1 never departed from wild-type dynamics, despite carrying the highest plaque density of the four recorded regions. Firing rate and spike-time irregularity changed in some vulnerable circuits and not others, and in both RSC and CA1, changes were present in young animals but absent in old. In the terms of our toy example, single-neuron statistics behaved like the amplitude of the oscillation and proximity to criticality behaved like its period. Interestingly, we required complementary analytical tools to capture disruptions related to criticality in the different vulnerable circuits. While all measures indicate the distance between observed data and true scale invariance, there are different ways by which scale invariance can manifest (and be measured). CA1 lost avalanche criticality (measured with DCC) in both amyloidosis and tauopathy. ACC lost purely temporal criticality (measured with *d*_2_), and RSC lost both. Strictly speaking, these are distinct forms of scale invariance, and it is possible that a network could produce one without the other^48^. Why disease may impact different forms of scale invariance in different circuits is an intriguing question.

In a brain comprising many connected neurons, patterns of activity differ wildly in terms of their size (number of involved neurons) and their duration. Intuitively, as pattern size increases, so does duration. When a brain changes states, the relationship that binds size and duration can also change. How states *can* alter interaction is not arbitrary — the scaling line defines the possible size/duration relationships that a brain can occupy. For example, elegant prior work demonstrates that, across behavioral state, species, and age, neural activity falls cleanly along the scaling line^51,52,54^. While this is a relatively nuanced concept from quasicriticality theory^52–54^, it is a powerful description of how neural activity can unfold within a connected brain. We used the scaling line to evaluate dynamics in wild-type and diseased animals. Position along the line, the feature that defines groups in all prior work, separated genotypes only in old RSC. To our surprise, *slope* separated them in a pattern that met all three of our criteria for a central, unifying feature of disease. Prior experimental and theoretical work illuminates a singular scaling line on which many systems fall. It is unknown whether and how the slope of the line itself can be modified. However, our data clearly reveal that, across molecular mechanisms and circuits, the slope of the scaling line is increased by pathology. Broadly, this should indicate that, in selectively vulnerable circuits faced with a molecular challenge related to neurodegeneration, activity patterns of a given size tend to take longer than one would expect in a healthy circuit. However, another measure of this relationship, the *γ* exponent, showed no significant change across conditions. These observations should push a small but growing theoretical field into new territory. It is not immediately clear how the rules governing the dissipation of information in a neuronal network are established. To elucidate a possible mechanism by which the scaling line slope might systematically change, we ran simulations, sweeping a range of parameters in a cortical branching model. We were able to recapitulate circuit-specific differences in activity and scaling line slope by increasing overall network strength, biasing each neuron’s synaptic weights towards a small subset of inputs, and disrupting intrinsic excitability.

If the loss of proximity to criticality contributes to cognitive decline, the dynamics of vulnerable circuits should be considered a viable target for intervention in addition to strategies that directly counteract the misfolded proteins that initiate disease^78–80^. Proximity to criticality is homeostatically regulated, and some of the mechanisms that implement this regulation are coming to light. After monocular deprivation, V1 returns to a near-critical regime before firing rates recover^41^, and modeling suggests that inhibitory interneurons are necessary for the homeostatic return^41,81^. Similarly, but under less severe perturbative conditions, sleep restores the near critical regime in isocortical networks after waking activity moves them away^42^. Taken together these observations suggest that, in the context of disease, careful control of inhibitory circuits and augmentation of some of sleep’s components may allow for the restoration of the dynamics upon which brain function depends. As a complement to strategies whose success is largely measured by reduced molecular burden, therapies directly focused on the restoration and/or maintenance of circuit function itself hold enormous promise.

## SUPPLEMENTAL FIGURES

**Figure S1:**
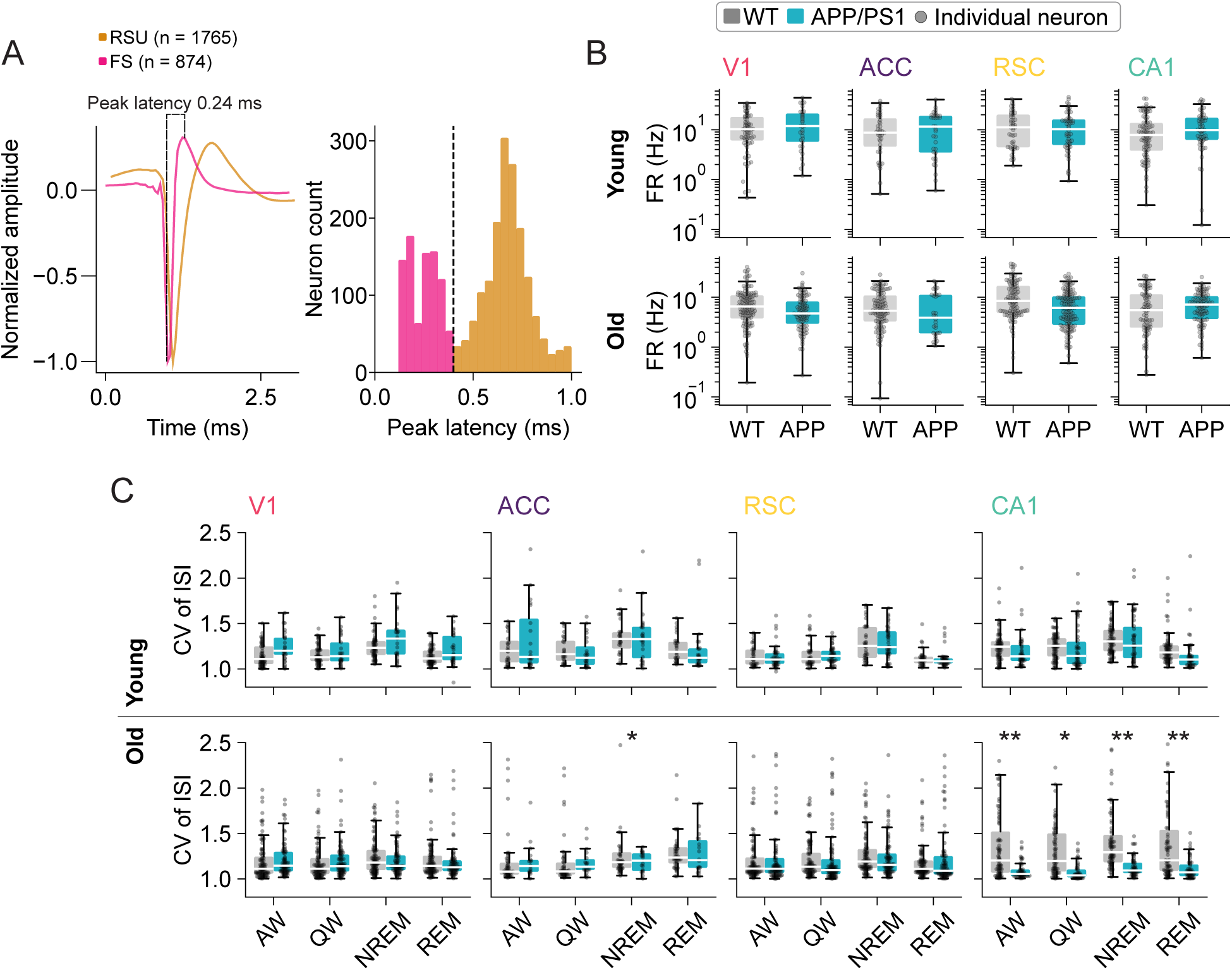
Cell-type classification and age-dependent changes in spike time variability in fast-spiking units. **A.** (left) Example mean waveforms of a representative regular-spiking unit (RSU, grey) and fast-spiking unit (FS, red), normalized to trough depth; peak latency (trough-to-peak duration) is the feature used to discriminate cell types. (right) Distribution of peak latency across all classified units, showing bimodal separation between FS (red) and RSU (grey) populations (RSU: n = 1765; FS: n = 874). Dashed line indicates the classification boundary. **B.** Mean firing rate of fast-spiking units as a function of genotype, region, and age group. Firing rate was computed per neuron across the entire recording epoch, collapsed across brain state (LMER: log(firing rate) ∼ genotype * age group * region * cell type + sex + prop_NREM_ + prop_REM_ + prop_active wake_ + prop_quiet wake_ + (1 animal), genotype contrasts FDR-corrected). No fast-spiking contrast reached significance. **C.** Coefficient of variation (CV) of the interspike interval for fast-spiking units as a function of genotype, region, age group, and brain state (* p *<* 0.05, ** p *<* 0.01; LMER: CV genotype * age group * region * cell type * state + sex + (1 animal/recording epoch/neuron), genotype contrasts FDR-corrected across state age group region cell type). APP/PS1 fast-spiking units showed reduced CV in old CA1 (all states) and old ACC (NREM). Grey dots indicate individual neurons.

**Figure S2:**
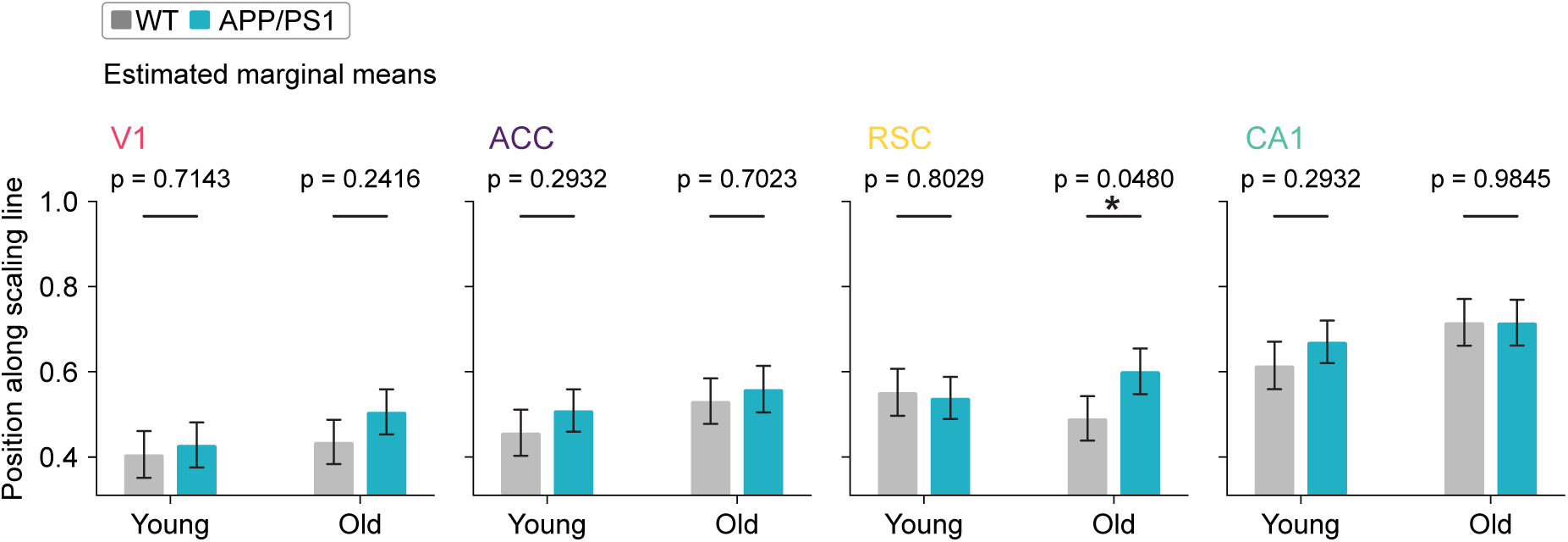
Position along the avalanche scaling line is largely preserved across genotypes. Position of each 2 h block along the pooled scaling line, as a function of genotype, age group, and brain region. For each region, a scaling line was fit to the (*τ_S_, τ_D_*) exponents of all blocks by total least squares, and each block’s position was computed as its projection onto the line, normalized to the unit interval across blocks; higher values indicate positions further along the scaling continuum. Bars indicate LMER-estimated marginal means SEM; circles indicate individual-animal median values. Genotype (APP/PS1 vs. WT) contrasts were computed within each age region combination (* p *<* 0.05, ** p *<* 0.01, *** p *<* 0.001; LMER: position genotype age group region + splines::ns(N, 3) + sex + (1 animal/recording epoch); genotype contrasts Benjamini–Hochberg FDR-corrected). Position differed significantly between genotypes only in RSC of aged animals (*p* = 0.048); all other comparisons were non-significant.

**Figure S3:**
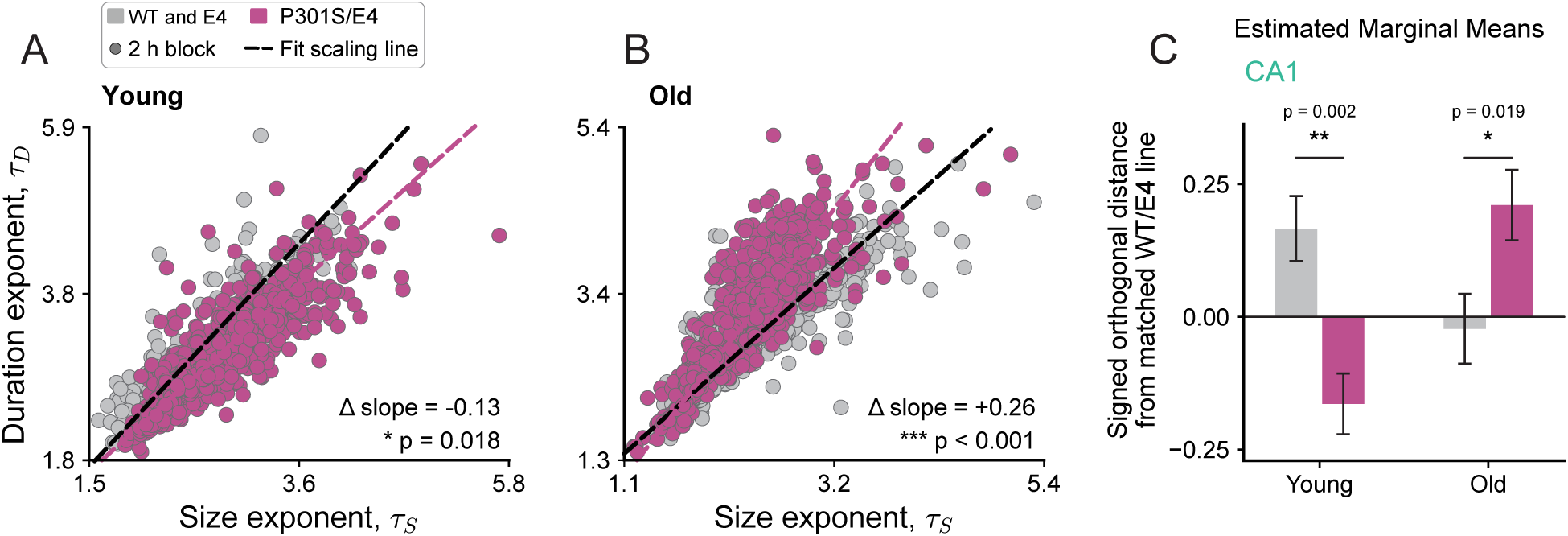
Departure from WT scaling line (unshifted) is present in CA1 in a mouse of tau-mediated neurodegeneration (P301S/E4). Same method as in Figure 5. **A.** Per-region *τ_S_*–*τ_D_* scatter on the scaling line from CA1 dynamics for young animals (3-4 mo). Each point is one 2 h block, colored by genotype (control [WT and E4] in grey, and P301S/E4 in magenta). **B.** Same as **A**, but for old animals (10-15 mo). **C.** Signed orthogonal distance of CA1 blocks from the control scaling line. For each age, a reference line was fit to control blocks alone by total least squares, and each block’s signed perpendicular distance from it was computed, so control blocks center near zero and positive values indicate displacement above the control line.

**Figure S4:**
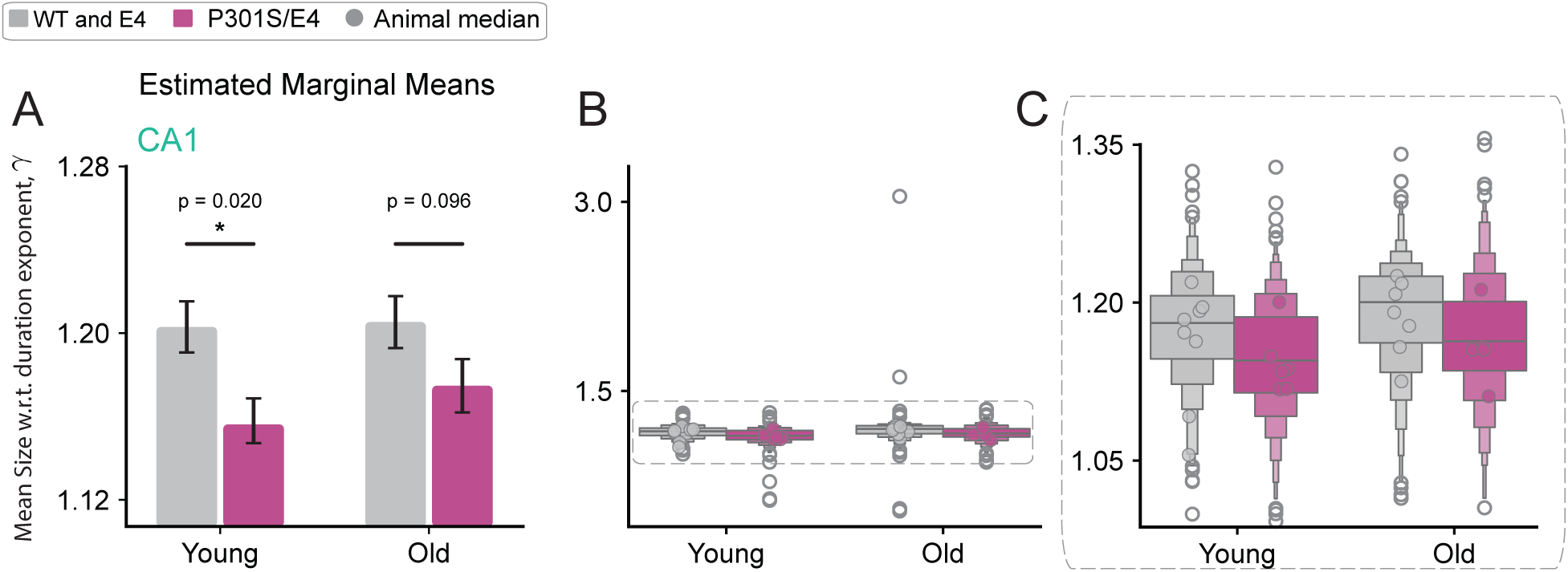
Differences between P301S/E4 and WT/E4 genotypes in *γ* exponents across ages. **A.** Estimated marginal means from LMER. Fitted *γ* exponents were fitted across WT/E4 and P301S/E4 genotypes showing significant differences for the young group (3-4 mo), but not for the old group (10-15 mo). (LMER: *γ* genotype age group + splines::ns(N, 3) + sex + (1 animal/recording epoch), Satterthwaite genotype contrasts and Benjamini–Hochberg FDR-corrected).**B.** Raw *γ* values. **C.** Raw *γ* values inset for central 99% of blocks.

**Figure S5:**
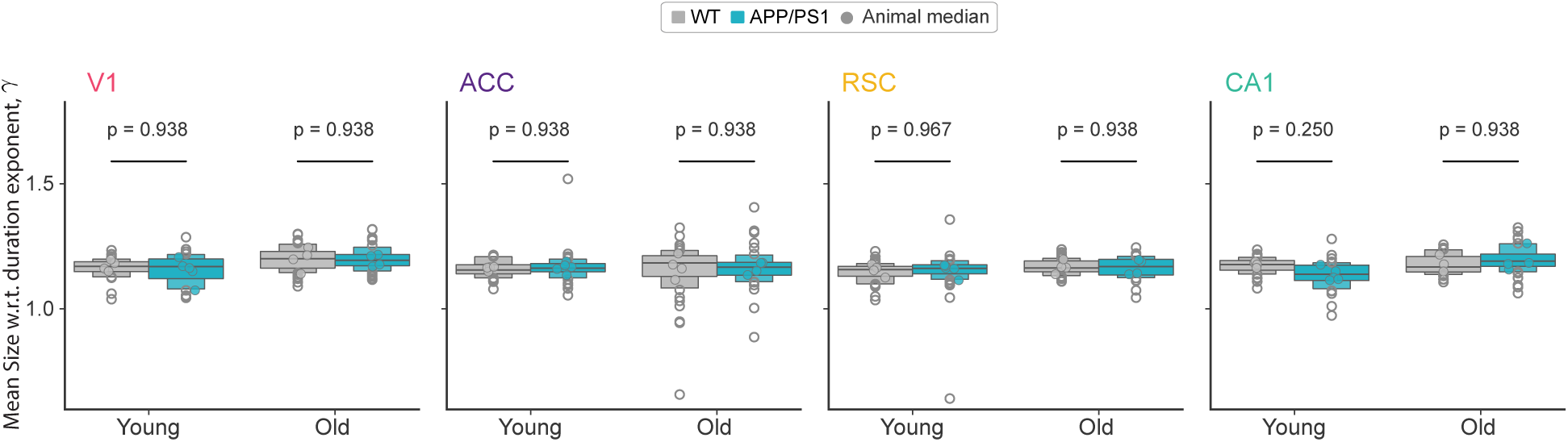
Differences between APP/PS1 and WT genotypes in *γ* exponents across ages and regions. Raw *γ* exponent values. (LMER: *γ* ∼ genotype ∗ age group ∗ region + splines::ns(N, 3) + sex + (1|animal/recording epoch), Satterthwaite genotype contrasts and Benjamini–Hochberg FDR-corrected).

**Figure S6:**
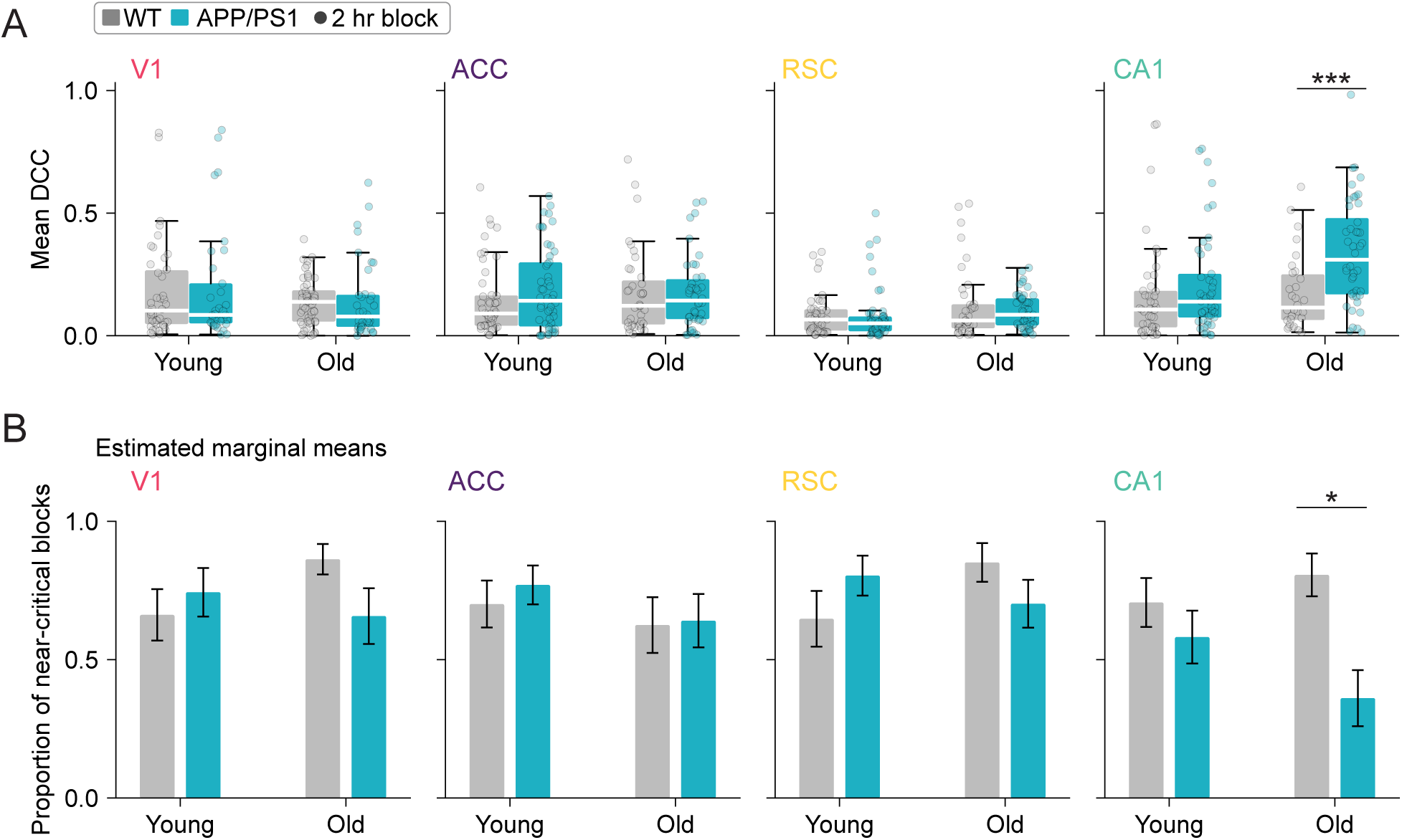
Age-dependent, circuit-specific disruption of near-critical dynamics in regular-spiking units in APP/PS1 mice. **A.** Mean deviation from criticality coefficient (DCC) across 2 h recording blocks, computed using regular-spiking units (RSUs) alone, as a function of genotype, age group, and brain region. The shaded yellow region denotes DCC 0.3 (near-critical). Each dot represents an individual 2 h block; boxplots show the distribution across blocks. (* p *<* 0.05, ** p *<* 0.01, *** p *<* 0.001; LMER: DCC genotype age group region + splines::ns(*n*_RSU_, 3) + sex + (1 animal/recording epoch); genotype contrasts Benjamini–Hochberg FDR-corrected.) **B.** Proportion of recording blocks classified as near-critical (DCC 0.3) based on RSU activity, as a function of genotype, age group, and brain region. Circles indicate the observed proportion of near-critical blocks for individual animals; bars and error bars indicate binomial GLMM-estimated proportions ± SEM. (* p *<* 0.05, ** p *<* 0.01, *** p *<* 0.001; binomial GLMM: near-critical genotype age group region + splines::ns(*n*_RSU_, 3) + sex + (1 animal:recording epoch); genotype contrasts Benjamini–Hochberg FDR-corrected.)

**Figure S7:**
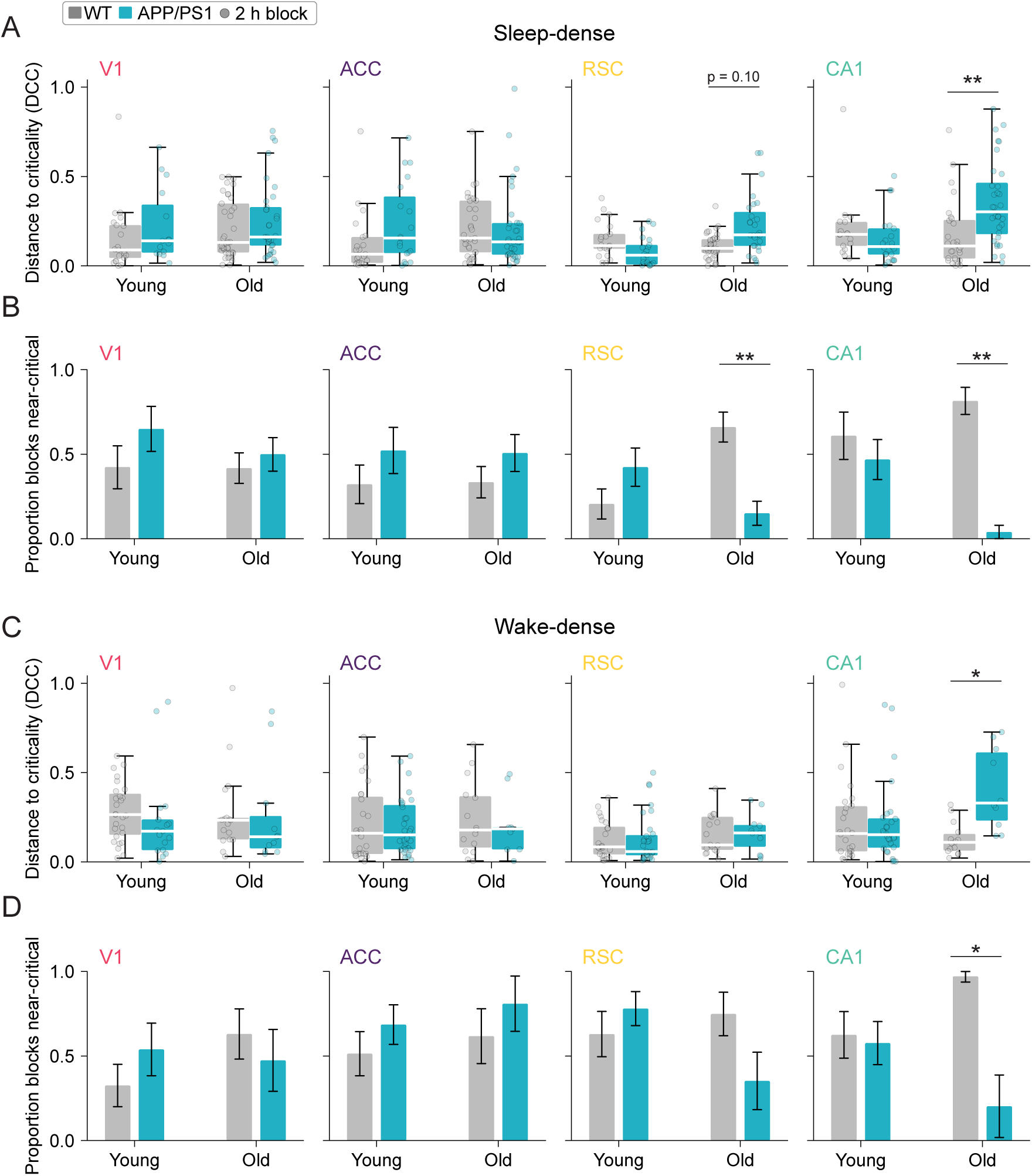
Near-critical neural dynamics are differentially disrupted in sleep- and wake-dense recording blocks in APP/PS1 mice. **A.** Mean deviation from criticality coefficient (DCC) across sleep-dense 2 h recording blocks as a function of genotype, age group, and brain region. Sleep-dense blocks were defined as blocks in which sleep comprised greater than 50% of the recording period. The shaded yellow region denotes DCC ≤ 0.3 (near-critical). Each dot represents an individual 2 h block; boxplots show the distribution across blocks. (* p *<* 0.05, ** p *<* 0.01, *** p *<* 0.001; LMER: DCC ∼ genotype * age group region + splines::ns(N, 3) + sex + (1 animal/recording epoch); genotype contrasts Benjamini–Hochberg FDR-corrected.) **B.** Proportion of sleep-dense recording blocks classified as near-critical (DCC ≤ 0.3) as a function of genotype, age group, and brain region. Circles indicate the observed proportion of near-critical blocks for individual animals; bars and error bars indicate binomial GLMM-estimated proportions SEM. (* p *<* 0.05, ** p *<* 0.01, *** p *<* 0.001; binomial GLMM: near-critical ∼ genotype * age group region + splines::ns(N, 3) + sex + (1 animal:recording epoch); genotype contrasts Benjamini–Hochberg FDR-corrected.) **C.** Same as in (**A**), but for wake-dense recording blocks, defined as blocks in which wake comprised greater than 50% of the recording period. **D.** Same as in (**B**), but for wake-dense recording blocks.

## METHODS

### Animals and datasets

All procedures were approved by the Washington University in St. Louis Institutional Animal Care and Use Committee, in accordance with the NIH Guide for the Care and Use of Laboratory Animals. Mice were housed in an enriched environment on a 12:12 h light:dark cycle with *ad libitum* food and water.

#### APP/PS1 recordings

We analyzed a previously published dataset^30^ of chronic, multi-region recordings from 17 mice of the APP/PS1 line (B6.Cg-Tg[APPswe,PSEN1dE9]85Dbo/Mmjax; MMRRC 034832-JAX^46^): 4 young wild-type (WT), 5 young APP/PS1, 4 old WT, and 4 old APP/PS1 mice. Young mice were 2–4 months of age, before substantial plaque accumulation, and old mice were 12–16 months of age, during advanced amyloidosis.

#### APP/PS1 amyloid plaque quantification

We previously quantified plaque density throughout cortex and hippocampus in a large independent cohort of APP/PS1 mice (3 mice at 4 months and 140 at 12–18 months) by serial two-photon tomography of methoxy-X04-labeled brains registered to the Allen Common Coordinate Framework^30,31^. We report those results without reanalysis.

#### P301S/E4 recordings

To compare neural activity features across mechanistically distinct diseases, we reanalyzed CA1 recordings from a previously published cohort^1^ of P301S/E4 mice, which express 1N4R human tau carrying the P301S mutation on a human APOE4 knock-in background and develop progressive tau-mediated neurodegeneration^82^, and controls. In this dataset, we recorded from 13 P301S/E4 mice, 23 WT mice, and 7 E4 control mice that express the APOE4 background but not the tau transgene and therefore do not develop tauopathy. Neural activity from single units in CA1 was recorded longitudinally at ages between P36 and P533. In the original study, WT and E4 mice did not differ in neuronal activity and were pooled as controls^1^. For the scaling line analysis, we considered 2 h recording blocks from mice belonging to young (90-120 d) or old (300-450 d) age groups. This analysis included 8 control and 6 P301S/E4 mice in the young group, and 7 control and 5 P301S/E4 mice in the old group. No animals contributed to both age groups.

### Surgery, electrophysiology, and spike sorting

Surgical implantation, electrophysiological recording, and spike sorting were performed as described previously^1,30,42,83^. Briefly, each APP/PS1 or WT mouse was implanted with four custom 64-channel nichrome tetrode arrays targeting V1, ACC, RSC, and CA1, and was recorded continuously in an enriched home cage for periods ranging from two weeks to three months (eCube, White Matter LLC). Neuronal signals were sampled at 25 kHz. Spikes were extracted and clustered with SpikeInterface^84^ and MountainSort4^85^, and single units were identified with a gradient-boosted decision tree trained on measures of cluster quality and isolation, followed by manual inspection^1^. The location of each array was confirmed post hoc by histological reconstruction. Approximately 24 h of continuous data were analyzed from each animal^30^.

### Cell-type classification

Single units were classified as regular-spiking (RSU, putative excitatory) or fast-spiking (FS, putative inhibitory) by the trough-to-peak duration of the mean spike waveform. Units with trough-to-peak durations greater than 0.4 ms were classified as RSU, and units with shorter durations as FS^39,86,87^. Trough-to-peak duration was distributed bimodally across populations of neurons and revealed that a 0.4 ms threshold cleanly separated the two classes (RSU, *n* = 1765; FS, *n* = 874; Fig. S1A).

### Sleep and arousal-state scoring

Arousal states were scored as described previously^30^. Briefly, LFP (0.1–60 Hz) from five manually selected channels per animal was averaged and used to generate spectrograms. Each 4 s epoch was manually assigned to active waking, quiet waking, NREM sleep, or REM sleep based on inspection of LFP spectrograms and consideration of delta power (high in NREM), theta power (high in REM), and movement activity (high in waking)^83,88,89^.

### Single-neuron firing rate and variability

Mean firing rate was computed for each single unit over the full clustering interval (typically 12 h), the timescale over which firing-rate homeostasis operates^38,39^, as the total spike count divided by the interval between the neuron’s first and last spike. Only units with a presence ratio greater than 0.99 were included (units whose spikes were distributed across more than 99% of the clustering interval). Therefore, for every included neuron, the interval between the first and last spike differs very little from the full clustering interval. Here, we used the interval between the first and last spike so that each firing rate is measured during the period over which the unit was demonstrably tracked. We also measured the coefficient of variation (CV) of the interspike interval (ISI) for each neuron. CV of ISI is defined as the standard deviation of the ISIs divided by their mean, computed separately for each neuron in each arousal state.

### Neuronal avalanches and the deviation from criticality coefficient (DCC)

Neuronal avalanches were measured as described previously^1,41^. Spike times from all included units (both RSU and FS) in a region were binned at 40 ms and summed to yield a population activity time series. Avalanches were defined as contiguous bins in which population activity exceeded the 35th percentile of its distribution. The size *S* of each avalanche was the total number of spikes above threshold, and its duration *T* was the number of bins it spanned. Within each 2 h block, the distributions of *S* and *T* were fit with truncated power laws, *P* (*S*) ∞ *S^−τS^* and *P* (*T*) *T ^−τD^*, by maximum likelihood^41,90^. Goodness of fit was assessed by comparing the Kolmogorov–Smirnov (KS) distance between each empirical distribution and its fit with the KS distances of 1000 synthetic power-law distributions with the same exponent, number of avalanches, and minimum and maximum values. A distribution passed if at least 5% of synthetic distributions had larger KS distances (*p* ≥ 0.05)^1^.

The exponent *γ* was fit by linear regression of log⟨*S*⟩ on log *T*, where ⟨*S*⟩(*T*) is the mean size of avalanches of duration *T*. The scaling relation predicts this exponent from the size and duration exponents as (*τ_D_*1)*/*(*τ_S_* 1)^41,60^. DCC was defined as the absolute difference between the fitted and predicted values of *γ*, such that values near zero indicate dynamics near criticality^1,41^. A block was classified as near-critical if both its size and duration distributions passed the goodness-of-fit test and its DCC was at most 0.3^1^.

For analyses restricted to regular-spiking units, avalanches, exponents, and DCC were recomputed from the summed activity of RSUs alone. For state-resolved avalanche analyses, blocks in which the animal spent more than 50% of the time asleep (NREM or REM) were classified as sleep-dense, and blocks in which it spent more than 50% of the time awake were classified as wake-dense.

### Scaling line analysis

The size and duration exponents of all blocks were analyzed in the *τ_S_*–*τ_D_* plane^51,52,54^. Scaling lines were fit by total least squares (TLS), which treats both exponents as measured with error, separately for each genotype within each region and age group, giving *τ_D_* = *m*^ *τ_S_* + *b*^^^.

#### Position along the common scaling line

For each region, a single line was fit by ordinary least squares to the (*τ_S_, τ_D_*) exponents of all 2 h recording blocks. We pooled genotypes and age groups to refer to both genotypes on one common axis. Duration exponents were shifted by the fitted intercept so that the scaling line passes through the origin. Each 2 h block was projected onto the unit vector along the line, giving a coordinate measured from the point at which the line crosses *τ_S_* = 0. Coordinates were divided by the largest value within that region, so position runs from 0 to 1, with larger values indicating blocks farther along the line toward larger exponents. Because the normalization is per region, positions are comparable between genotypes and ages within a region but not across regions. Genotypes were compared with a linear mixed-effects model, position ∼ genotype ∗ age group ∗ region + splines::ns(*N*, 3) + sex + (1|animal/recording epoch), fit by REML with the bobyqa optimizer, using Satterthwaite degrees of freedom and Benjamini–Hochberg FDR-corrected genotype contrasts within each age × region combination (751 blocks, 35 recording epochs, 17 animals) (Fig. S2).

#### Slope of the scaling line

We assessed significant differences between genotypes in the slopes of the scaling lines with a linear mixed-effects model: *τ_D_*∼ *τ_S_* ∗ genotype ∗ age group ∗ region + splines::ns(*N,* 3) + sex + (1 animal/recording epoch) (Statistical analysis). Genotype differences in the *τ_S_* slope were estimated within each age group and region with emtrends^91^, using Kenward–Roger degrees of freedom and Benjamini–Hochberg FDR correction. This mixed-model slope regresses *τ_D_* on *τ_S_* and is a different estimator from the total least-squares slope used to draw and align the scaling lines: the dashed lines in each panel are the per-genotype TLS fits, whereas the Δslope and significance annotated on the panel are the emtrends genotype contrast and its FDR-corrected *p*-value. The two estimators need not agree numerically.

#### Signed distance from the wild-type line

Within each region and age group, a reference line was fit by TLS to WT blocks alone. The signed orthogonal distance of each block from this line was computed as the perpendicular distance between its (*τ_S_, τ_D_*) coordinate and the line, with positive values indicating a block above the line. By construction, WT blocks are centered near zero.

#### Alignment at (1, 1)

Any line that satisfies the scaling relation *τ_D_* 1 = *γ*(*τ_S_* 1) passes through (1, 1)^60^. Measured lines are offset from this point by thresholding, finite recording time, and error in the fitted exponents. Following prior work^52^, the duration exponents of each genotype within each region and age group were translated so that its TLS line passed through (1, 1),

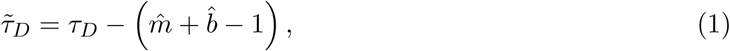

which leaves each slope unchanged. In this frame the WT line is *τ̃_D_* = *m*^ _WT_ *τ_S_*+ (1 − *m*^ _WT_), and the signed orthogonal distance of block *i* from it is

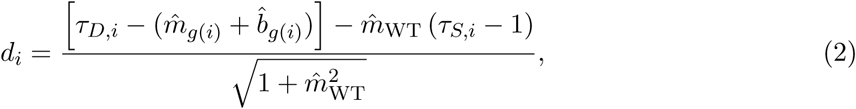

where *g*(*i*) is the genotype of block *i*. Because both lines pass through (1, 1), a given difference in slope produces larger distances in blocks farther from that point.

#### Scaling line analysis in P301S/E4 tauopathy dataset

The same avalanche scaling line analysis was applied to single-unit recordings of CA1 from P301S/E4 and control mice published in a prior paper^1^. Slopes of scaling lines were compared with linear mixed effects: *τ_D_* ∼ *τ_S_* ∗ genotype ∗ age group + splines::ns(*N,* 3) + sex + (1|animal/recording epoch). Because only CA1 was recorded in this dataset, the LMER models omit region. genotype differences in the *τ_S_* slope were estimated within each age group with emtrends^91^ using Kenward–Roger degrees of freedom. Signed distances were compared with signed distance ∼ genotype ∗ age group + splines::ns(*N,* 3) + sex + (1|animal/recording epoch), with Satterthwaite genotype contrasts within each age group. Both outcomes were Benjamini–Hochberg FDR corrected across the two age groups. Non-independence among 2 h blocks recorded from the same animal and recording epoch is absorbed by the nested random intercepts, and under the age windows used here no animal contributed blocks to both age groups.

### Distance to criticality (*d*_2_)

To measure proximity to criticality from the temporal structure of population activity rather than from avalanches, we computed *d*_2_^35,48^. For each 3-minute recording window, spikes from all included units in a region were binned at 15 ms and summed across the population, and the resulting time series was z-scored. An autoregressive model of order *p* = 20 was fit to this series by solving the Yule–Walker equations, yielding coefficients *φ*_1_*, … , φ_p_*, and

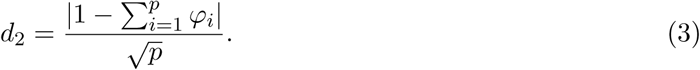

The sum Σ*_i_φ_i_* approaches 1 as population activity approaches a marginally stable, critical regime, so *d*_2_ approaches zero at criticality and increases toward 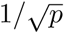 as temporal structure is lost, as in white noise. For the analysis of wake and sleep, only windows spent entirely in wake (active or quiet waking) or entirely in sleep (NREM or REM) were included.

### Computational model

We simulated the cortical branching model (CBM)^53,54^, a probabilistic cellular automaton. Each network comprised *N* = 128 randomly and strongly connected nodes, each receiving input from *k*_in_ = 10 neighbors. The probability that node *i* is activated by node *j* depends on the rank *n_ij_* ∈ {1*, … , k*_in_} of that connection among the inputs to node *i*,

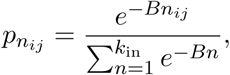

so that at *B* = 0 all inputs are equally effective, and as *B* increases, activation depends increasingly on the strongest inputs. The strength of each interaction is *P_ij_* = *κ p_nij_*, where the control parameter *κ* tunes the network toward the peak of susceptibility. Nodes were updated synchronously in discrete time. An active node activated each quiescent target independently with probability *P_ij_*. After activation, a node could not be reactivated for its refractory period, which was fixed at *τ_r_* = 1, so a node could be activated again on the step after it was active. Spontaneous activity was introduced as single-node activations at Poisson-distributed intervals with mean 1*/*(*p_s_N*) time steps, where *p_s_* is the probability of spontaneous activation.

The reference model, representing WT mice, had *κ* = 1.0062, *B* = 1.5, and *p_s_* = 3.16 × 10*^−^*^4^. These parameters place the reference network slightly below criticality, which maximizes the number of decades spanned by the avalanche distributions and avoids the bump in the distributions that finite network size produces near criticality, so that exponents can be fit cleanly. Each scaling line comprised 100 simulations, cycling through 6 independent random networks. In each simulation, *p_s_* and *κ* were drawn uniformly within ±20% and ±2% of the network’s values, which moves the network along its scaling line, and 3000 avalanches were recorded. Avalanche sizes and durations from each simulation were fitted with a discrete power law truncated to the best window [*x*_min_*, x*_max_] that captures scale invariance, where *x* is either avalanche size or duration. For a given window the exponent was obtained by maximum likelihood. *x*_max_ was set to the 99th percentile of that simulation’s sizes (or durations), and *x*_min_ was then chosen as the candidate minimizing the Kolmogorov–Smirnov distance between the fitted and empirical distributions^90^. Each window was required to span at least 1.5 decades for sizes and 1.0 decade for durations and to contain at least 500 avalanches.

To identify parameter changes sufficient to reproduce the region-specific changes in slope, we swept *κ*, *p_s_*, and *B*, with *k*_in_ and *τ_r_* fixed, over

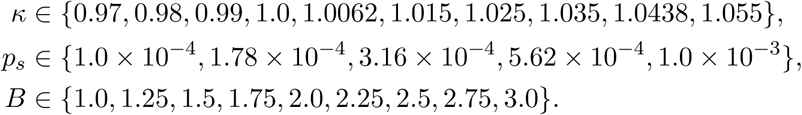

For each region, we compared four differences between groups: the mean, variance, CV of population activity, and the slope of the scaling line. For the data, these differences were APP/PS1 — WT in old animals, with population activity taken from the thresholded time series used to detect avalanches. For the model, they were region model − reference. Because the model and the data differ in absolute scale, parameter combinations were selected first by matching the sign of all four differences and then by matching their approximate magnitude. For display, each bar difference *v* was rescaled as sign(*v*) 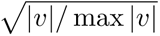 to lie in the interval [-1,1] (Fig. 8B). RSC and CA1 matched in the sign of all four differences in the data, so they converged on the same parameters.

### Statistical analysis

Statistical analyses used linear mixed-effects models (LMER) fit with lme4^92^ and lmerTest^93^ or, for binary outcomes, binomial generalized linear mixed models (GLMMs) fit with lme4, in R. Because neurons, epochs, and recording blocks are nested within animals, animal was the unit of replication, and the random effects of every model grouped observations by animal. Estimated marginal means and genotype contrasts (APP/PS1 vs. WT, or P301S/E4 vs. control) were computed with emmeans^91^ within each age × region combination, or age × region × state where applicable. Within each analysis, raw *p*-values were corrected across all genotype contrasts with the Benjamini–Hochberg procedure^94^ (for example, across the 16 state × age × region contrasts for *d*2), and corrected values are reported as *p*.

#### Single-neuron activity measures

The log firing rate of each neuron was modeled with genotype, region, age group, and cell type as fully interacting fixed effects, sex and the proportion of the clustering interval spent in NREM, REM, active waking, and quiet waking as covariates, and animal as a random intercept: log(firing rate) ∼ genotype×region×age group×cell type+sex+prop_NREM_+ prop_REM_ + prop_AW_ + prop_QW_ + (1 | animal). Because firing rate was measured once per neuron, animal was the only grouping factor. The CV of the ISI, resolved by arousal state, was modeled as CV ∼ genotype × age group × region × cell type × state+sex+(1 | animal/recording epoch/neuron). RSU and FS results from these models are shown separately (Fig. 1; Fig. S1).

#### Population activity measures

DCC, *γ*, signed distance from the WT line (unaligned and aligned), and *d*_2_ were modeled at the level of the block or window with genotype, age group, region, and, for *d*_2_, arousal state as fully interacting fixed effects, sex and the number of recorded units as covariates, and recording epoch nested within animal as random intercepts: *y* ∼ genotype × age group × region (× state) + splines::ns(*N,* 3) + sex + (1 | animal/recording epoch). The number of units *N* was entered as a natural cubic spline because avalanche statistics depend nonlinearly on the number of recorded neurons. For analyses restricted to RSUs, the spline was computed on the number of RSUs. These models were fit by restricted maximum likelihood, and degrees of freedom were estimated with Satterthwaite’s approximation.

#### Scaling line slope

### Slopes were compared with the model

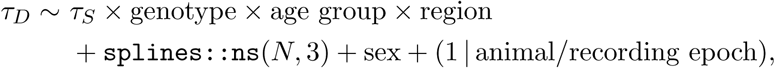

and genotype differences in the *τ_S_* slope were extracted within each age group and region with emtrends.

#### Proportion of time near-critical

Binarized near-critical status (yes or no) was modeled with a binomial GLMM of the same fixed-effect structure as the population measures, fit by maximum likelihood with the Laplace approximation. The full nested random-effects structure yielded a singular fit, with the variance among animals estimated at zero, so these models used a single grouping factor of recording epoch within animal, (1 | animal:recording epoch). The model estimates the proportion of near-critical blocks in each genotype, age group, and region, and genotype contrasts are reported as odds ratios.

### Sex as a biological variable

Sex differences in amyloid progression and disease severity are established in this model^95,96^. Thus, sex was included as a fixed-effect covariate in all mixed-effects models. All genotype effects therefore account for sex. Our study is underpowered to test specific hypotheses regarding sex differences.

## Data Availability

Data for the first paper based on this dataset^30^ are posted publicly (DOI 10.6084/m9.figshare.33011039). The raw broadband data are ∼ 100 terabytes and therefore cannot be easily shared publicly, but are available upon reasonable request.

## Code Availability

Code crucial to the electrophysiological recordings and data processing (e.g., spike sorting) is publicly available at github.com/hengenlab. Code used for the specific analyses in this paper are available at *github.com/hengenlab/APPPS*1 *and tau disrupt near critical dynamics*.

## Acknowledgments

This work is supported by NIH R01NS118442 and R01AG091773 (KBH), Cure Alzheimer’s Fund (JNM & KBH), Brain & Behavior Research Foundation Young Investigator Grant 29225 and in part by the National Institute on Aging R01AG047589 (JAH), NIH NINDS F32NS138250 and Center for Theoretical and Computational Neuroscience at WUSTL (LJF). We thank Kiran Bhaskaran-Nair, Gemechu Tolossa, and Sam Sooter for assistance with the development of code and use of analytical methods and David Borchelt for sharing APP/PS1 mice.

## Author Contributions

J.N.M.: Conceptualization, Methodology, Investigation, Formal analysis, Visualization, Writing – original draft, Writing – review & editing. L.J.F.: Formal analysis, Methodology, Investigation, Visualization, Writing – original draft, Writing – review & editing. S.J.B.: Conceptualization, Methodology, Investigation, Data curation, Writing – original draft, Writing – review & editing. C.F.: Investigation, Data curation. B.T.H.: Investigation, Data curation. Z.B.R.: Investigation, Data curation. H.E.: Investigation, Formal analysis. K.K.: Formal analysis. J.D.W.: Conceptualization, Methodology, Supervision. J.A.H.: Conceptualization, Methodology, Supervision. W.L.S.: Conceptualization, Methodology, Formal analysis, Writing – review & editing. D.M.H.: Conceptualization, Methodology, Supervision, Writing – review & editing. K.B.H.: Conceptualization, Methodology, Supervision, Project administration, Visualization, Writing – original draft, Writing – review & editing.

